# A STAT5B-driven mouse model of hepatosplenic γδ T-cell lymphoma reveals therapeutic efficacy of JAK inhibition

**DOI:** 10.1101/2025.10.24.684333

**Authors:** Myint Myat Khine Aung, Susann Schönefeldt, Sophie Pfalz-Kraupp, Tamara Wais, Tobias Suske, Safia Zahma, Stefan Franz, Andrea Müllebner, Gabriela Feurstein, Christina Wagner, Thomas Eder, Tea Pemovska, Christiane Agreiter, Alexander Pichler, Philipp B Staber, Martin Hofer, Ralf Steinborn, Svenja-Verena Class, Ingrid Simonitsch-Klupp, Marcus Bauer, Andreas Wilfer, Thomas Weber, Swapnil Potdar, Tero Aittokallio, Dennis Jungherz, Tony A Müller, Julia List, Dagmar Gotthardt, Florian Grebien, Vasileios Bekiaris, Marco Herling, Richard Moriggl, Heidi A Neubauer

## Abstract

Hepatosplenic T-cell lymphoma (HSTCL) is a rare and aggressive neoplasm associated with poor responses to standard chemotherapy regimens and low survival rates. No targeted therapies are available for HSTCL, and preclinical models to test new treatment options have not been established. The JAK-STAT signaling cascade is a key dysregulated pathway in HSTCL, and *STAT5B^N642H^*is the most frequent somatic mutation in the disease. Here, we report on newly established clonal, murine γδ T-cell lymphoma cell lines initiated and driven by oncogenic *STAT5B^N642H^*, which recapitulate key immunophenotypic features, gene expression profiles and typically low cytolytic activity of patient-derived human HSTCL cells. CRISPR-Cas9 mediated knockout demonstrated growth dependence on *STAT5B^N642H^*. Murine C15 cells were allo-engrafted intravenously into both immunodeficient and immunocompetent mice to model an aggressive HSTCL-like disease at high penetrance, with recipient mice displaying hepatosplenomegaly and destructive γδ T cell organ infiltration, including bone marrow and blood involvement. We identified the potential of JAK inhibition as a targeted treatment strategy for HSTCL, and found the clinically approved JAK inhibitor upadacitinib to display selective anti-tumor efficacy against *STAT5B*-mutated HSTCL cell lines *in vitro, in vivo*, and in primary HSTCL patient samples. Overall, we describe the first robust STAT5B-driven preclinical model resembling features of HSTCL in an immune competent setting. This tool is expected to accelerate the study of HSTCL disease mechanisms and the testing of novel therapies. Our data further present the JAK inhibitor upadacitinib as a promising targeted treatment option for *STAT5B*-mutated HSTCL.

## Introduction

Hepatosplenic γδ T-cell lymphoma (HSTCL) is a rare, aggressive mature T-cell lymphoma that manifests predominantly in young adults with a median age of 34 years.^1,2^ HSTCL is characterized by the expansion of malignant γδ T cells primarily in the liver and spleen, causing hepatosplenomegaly. Tumor cell infiltration into the bone marrow (BM) and involvement of peripheral blood (PB) is also often observed.^2^ HSTCL typically shows a rapidly progressing course and an overall poor response to treatments.^3,4^ Preferred first-line therapeutic strategies for HSTCL patients include platinum/ifosfamide-based induction regimens followed, when feasible, by allogeneic hematopoietic stem cell transplantation.^3^ There are no targeted therapies approved for HSTCL and patients face a dismal prognosis with a five-year overall survival rate of 7% and a median survival of 13 months.^5,6^ The rarity of the disease limits the study of new therapeutic options using patient-derived tumor samples or clinical trials, and there are currently no established preclinical models available. Hence, there is an urgent need for faithful models to facilitate the preclinical testing of new targeted therapies for HSTCL.

Genetic studies have reported specific cytogenetic aberrations in HSTCL patients, the most common being isochromosome 7q, occurring in up to 70% of cases, and trisomy 8, in up to 50% of patients.^2,7^ Key pathways frequently dysregulated in HSTCL include JAK-STAT, PI3K-AKT-mTOR and epigenetic regulation.^6,8^ Specifically, loss-of-function (LOF) mutations were reported in *SETD2*, *INO80*, *TET3* and *SMARCA2*, whereas the most common gain-of-function (GOF) mutations occur in *STAT5B*, *STAT3* and *PIK3CD*.^9–11^ Notably, around one-third of HSTCL patients harbor the oncogenic *STAT5B^N642H^*mutation, representing the most frequent mutation in this disease.^8^ Patients with T-cell malignancies harboring the *STAT5B^N642H^* mutation were reported to have poorer clinical outcomes and a higher risk of relapse compared with *STAT5B* wildtype (WT) patients.^12,13^ We previously demonstrated that the *STAT5B^N642H^*oncogene efficiently induces γδ T-cell transformation in mice.^14^ Therefore, we reasoned that establishing a robust, cell line-based γδ T-cell lymphoma (γδTCL) allograft mouse model driven by STAT5B^N642H^ could recapitulate features of HSTCL and could facilitate the preclinical testing of new therapeutic options.

Here, we have generated clonal murine γδTCL cell lines dependent on oncogenic STAT5B^N642H^, which display immunophenotypic, transcriptional and functional features of human HSTCL cells. The C15 cell line can be engrafted intravenously into immunodeficient and immunocompetent mice, generating the first preclinical *in vivo* model with aggressive, highly penetrant γδ HSTCL-like disease. Notably, the JAK inhibitor (JAKi) upadacitinib displayed significant and selective anti-tumor efficacy against *STAT5B*-mutated HSTCL cells *in vitro* and *in vivo*, and in primary HSTCL patient samples, highlighting upadacitinib as a potential targeted therapeutic option for *STAT5B*-mutated HSTCL.

## Methods

### Study approvals

All mouse breeding and experiments were authorized by the Austrian Federal Ministry of Education, Science and Research (BMBWF-68.205/0084-V/3b/2019, 2022-0.118.098, 2023-0.651.094, 2024-0.191.292, BMWFW-68.205/0093-WF/V/3b/2015 and 2022-0.404.452).

Collection and use of healthy-donor derived T cells isolated from buffy coats for RNA-seq were approved for research purposes by the ethics committees of the University Hospital of Cologne (#19-1089).

Viable, cryopreserved samples from the Viennese Viable Biobank for Heamatological Diseases (Vivibank, EK 1284/18) and associated FFPE tumor samples from HSTCL patients were obtained with approval from the ethics committees of the Medical University of Vienna (EK 1448/2024). The research was conducted in accordance with the Declaration of Helsinki.

### Generation of murine clonal cell lines

Wildtype C57BL/6N mice transplanted with γδ T cells isolated from lymph nodes of *Vav1*-hSTAT5B^N642H^(FLAG) transgenic mice developed a γδ T-cell lymphoma/leukemia phenotype, as previously reported (a brief description of the transgenic mouse model is given in the **Supplemental Methods**).^14^ Cryopreserved lymph node cells from one terminally diseased animal were thawed and 2×10^6^ cells were incubated at 37°C in 5 ml media containing RPMI 1640 (Gibco) with 20% heat-inactivated fetal bovine serum (hi-FBS; Biowest), 2mM L-glutamine (Gibco), 10 U/ml penicillin/streptomycin (Biowest), 55 μM β-mercaptoethanol (Gibco) and 40 ng/ml recombinant human interleukin-2 (IL-2; ImmunoTools GmbH). Every seven days, 2.5 ml of the media was removed and replaced with 2.5 ml fresh media containing 110 μM β-mercaptoethanol and 80 ng/ml IL-2. After two weeks, large, proliferative cells with mostly adherent properties were observed and expanded as detailed below. Clonal cell lines were further generated by FACS sorting single-cells into a 96-well plate using a FACSAria III cell sorter (BD Biosciences) and expanding in standard media with 10 ng/ml IL-2 and Normocin^TM^ (Invitrogen). The clonal cell lines were cultured in the presence of Normocin^TM^ until they had been expanded into 25 cm² cell culture flasks, to minimise the risk of bacterial contamination.

### Cell line drug treatments

Drugs were purchased from MedChemExpress and reconstituted in DMSO. Cells were seeded in triplicates into flat-bottom 96-well plates (Greiner AG) at 2×10^4^ cells/well. The following day, cells were treated with serial two-fold or five-fold dilutions of the drugs, or DMSO as a negative control, in 10 ng/ml IL-2 supplemented media. Bortezomib (100 μM) served as a positive control. Cells were incubated at 37°C for 48 hr. Cell viability was measured using a CellTiter-Blue Cell Viability Assay (Promega) on a GloMax Discover Microplate Reader (Promega). IC_50_ values were determined using Breeze software.^15^

### Drug sensitivity profiling in primary patient cells

Spleen, BM or PB samples from HSTCL patients (**Supplemental Table 1**) were cryopreserved after Ficoll-Paque mononuclear cell separation. Cell viability profiling was performed as previously described.^16^ After 24 hr drug treatment, cells were stained with DAPI and antibodies marking tumor or healthy cells, and multiplex cell analysis was performed using high-throughput flow cytometry.

### *In vivo* transplant models and JAK inhibitor treatment

M2, C15 and DERL-7 cells were stimulated with 10 ng/ml IL-2 two hr prior to transplantation. M2 or C15 cells (1×10^6^ in 100 µl PBS) were intravenously administered to C57BL/6 Ly5.1 or NSG mice via tail vein injection. For *in vivo* upadacitinib treatment, 10 days post transplantation C15-recipient mice were randomly distributed into two groups and treatment commenced of once-daily oral gavage with 10 mg/kg upadacitinib or vehicle, five days per week in two treatment cycles. DERL-7 cells (2×10^6^ in 100 µl PBS) were intravenously administered to NSGS mice via tail vein injection. Further details of the transplant models, drug treatment and downstream analyses are given in the **Supplemental Methods**.

## Results

### Murine γδ T cell lines harboring the *STAT5B^N642H^* mutation display HSTCL-like features

Primary murine γδ T cells expressing human STAT5B^N642H^ induce an aggressive γδTCL when transplanted from transgenic mice into wildtype (WT) C57BL/6 mice.^14^ However, STAT5B^N642H^ transgenic mice are challenging to maintain due to the rapid onset of aggressive CD8^+^ T cell disease.^17^ Thus, we sought to establish STAT5B^N642H^ expressing murine γδTCL cell lines as robust models for *in vitro* and *in vivo* studies. Using *ex vivo* tumor cell outgrowth cultures from *STAT5B*-mutant γδ T cell recipient mice, we established polyclonal γδTCL cell lines in the presence of IL-2 (**Figure 1A**). To recapitulate the clonal nature of HSTCL, we subsequently performed single cell sorting to generate clonal lines. These clones were screened by immunoblotting for FLAG-tag expression (indicating the presence of STAT5B^N642H^ protein) and strong STAT5 activation (phospho-tyrosine 694/699; pY-STAT5) (**Supplemental Figure 1A**). Three clonal lines, C2, C6 and C15, were selected and the expression of the STAT5B^N642H^ transgenic protein was further confirmed by flow cytometric analysis of FLAG expression and by Sanger sequencing (**Supplemental Figure 1B-C**). In culture, these lines displayed largely adherent properties and were morphologically distinct (**Supplemental Figure 1D**).

**Figure 1:**
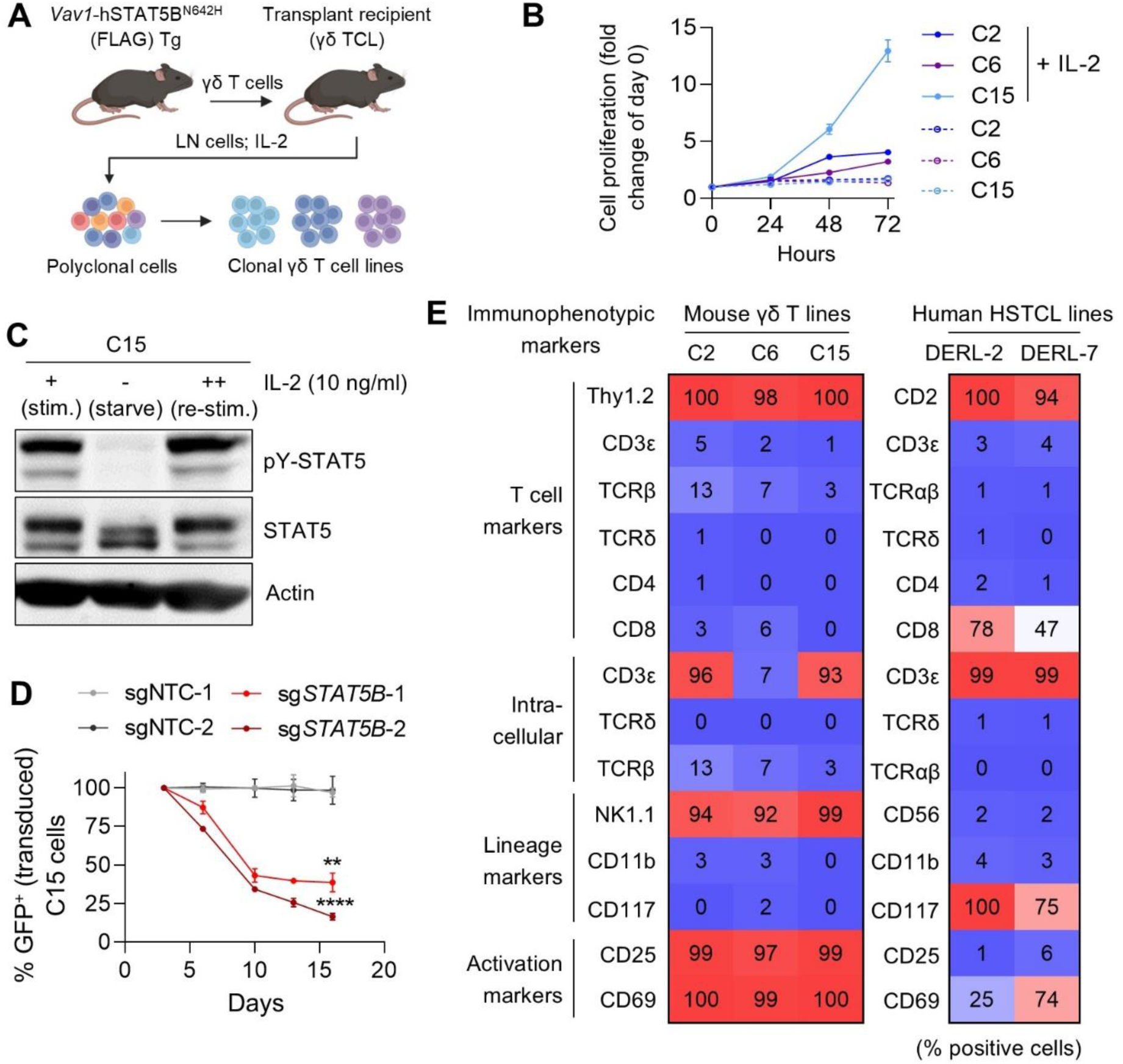
Generation of clonal murine STAT5B^N642H^-expressing γδ T-cell lymphoma cell lines with HSTCL-like features. **A)** Schematic depicting the procedure taken to generate clonal murine γδ T-cell lines from a human (h)STAT5B^N642H^ transgenic (Tg) mouse.^17^ LN, lymph node; TCL, T-cell lymphoma. **B)** Cell proliferation of murine clonal C2, C6 and C15 γδ TCL cell lines in the presence or absence of IL-2 over 72 hr, measured by flow cytometry. Data are graphed as mean (± SD) of technical triplicates from one experiment, representative of three independent experiments (*n* = 3). **C)** Western blot showing STAT5 activity in C15 cells cultured in media supplemented with IL-2 for 24 hr (+), starved of IL-2 for 8 hr (-), and then restimulated with IL-2 for 1 hr (++). Immunoblotting for pY-STAT5 and total STAT5 was performed, with actin serving as a loading control (blots representative of three independent experiments; *n* = 3). **D)** Cell competition assay of C15 cells transduced with human *STAT5B*-targeting sgRNAs or non-targeting control (NTC) sgRNAs, both containing GFP markers, for CRISPR/Cas9 gene editing. Percentages of GFP+ cells were measured over time using flow cytometry. Data are graphed as mean (± SD) of technical triplicates from one experiment, representative of three independent experiments (*n* = 3). \*\**p* < 0.01, \*\*\*\**p* < 0.0001 (compared with sgNTC-1); two-way ANOVA with Tukey post-test for multiple comparisons. **E)** Heatmap displaying percentage expression of various cell surface and intracellular protein markers from murine C2, C6 and C15 cell lines, as well as from human DERL-2 and DERL-7 HSTCL cell lines, measured by flow cytometry. Data are displayed as average values from three independent experiments (*n* = 3).

To further profile our cell lines, we performed detailed biochemical and immunophenotypic characterization with side-by-side comparisons of the two available human HSTCL cell lines, DERL-2 and DERL-7, established from a single HSTCL patient^18^ and harboring the *STAT5B^N642H^* mutation.^19,20^ Sanger sequencing revealed that the DERL-2 cell line is heterozygous for *STAT5B^N642H^*whereas the DERL-7 cell line displays *STAT5B^N642H^* homozygosity (**Supplemental Figure 2A**). The murine cell lines were dependent on IL-2 for sustained proliferation, with no cell proliferation observed in the absence of IL-2 across all three lines (**Figure 1B**). This was also the case for the human HSTCL cell lines, as shown (**Supplemental Figure 2B**) and previously reported.^18^ The C15 line showed faster proliferative kinetics, similar to the human DERL cells, as compared to the C2 and C6 lines. Despite the N642H mutation causing structural changes in STAT5B leading to stabilized and prolonged tyrosine 699 phosphorylation,^14^ STAT5 activation in the murine and human cell lines remained dependent on IL-2 stimulation as shown by the almost complete loss of pY-STAT5 signal upon IL-2 withdrawal, which was restored upon IL-2 re-stimulation (**Figure 1C**, **Supplemental Figure 2C-D**). IL-2 withdrawal was paralleled by loss of proliferative capacity, indicating a dependence of these cell lines on sustained IL-2-mediated oncogenic STAT5B activity. To confirm dependence on the driver STAT5B mutant, we performed a cell competition assay using C15 cells. We induced CRISPR-Cas9 mediated knockout of the *STAT5B^N642H^* transgene, without affecting endogenous *Stat5b*, and measured cell survival over 21 days. Indeed, loss of the STAT5B^N642H^ mutant resulted in significant depletion of the cells over time, in contrast to cells expressing a non-targeting negative control single guide RNA (sgRNA) (**Figure 1D**).

The immunophenotypic profiles of the murine cell lines showed strong similarities with the profiles of the human HSTCL cells, as determined by flow cytometry using a comprehensive set of surface and intracellular markers. Shared features included positive surface expression of pan-T cell marker Thy1.2/CD2 and the activation marker CD69 (highly expressed on DERL-7 cells), an absence of surface CD3ε, T-cell receptor (TCR)δ, TCRβ and CD4, and strong intracellular CD3ε staining (**Figure 1E**, **Supplemental Figure 3A-D**). All cell lines were negative for intracellular TCRδ and TCRβ, as well as the lineage marker CD11b, with positive surface CD117 expression detected on the human HSTCL cell lines indicating a more progenitor-like tumor cell of origin. While a lack of surface CD3 and γδTCR is uncharacteristic of primary HSTCL tumor cells,^2^ this is in line with the previously reported immunophenotype of the DERL cell lines.^18^ HSTCL tumor cells are typically CD8^-^, although some cases were reported to express CD8;^2^ the DERL cell lines displayed variable CD8 expression and the murine cell lines were consistently CD8^-^. Surprisingly, in our hands the DERL cell lines were CD56^-^, despite being derived from a patient with CD56^+^ disease.^18^ The murine cell lines expressed the NK-cell marker NK1.1 and additionally displayed consistent CD25 (IL2Rα) surface expression, potentially due to the fact that *Il2ra* is a bona fide STAT5 target gene^21,22^ (**Figure 1E**, **Supplemental Figure 3A-D**). The absence of intracellular FOXP3 protein ruled out a Treg phenotype (**Supplemental Figure 3E**).

Despite the absence of detectable TCRδ protein by flow cytometry, TCR rearrangement analysis by PCR confirmed rearrangements of both Vγ2 and Vγ4 as well as Vδ4 (nomenclature based on the Heilig and Tonegawa’s system^23^) in all three murine lines (**Supplemental Figure 3F**).

### STAT5B^N642H^ drives HSTCL-like transcriptional profiles in murine γδ T cells

To gain insights into the contribution of the STAT5B^N642H^ mutant transcription factor towards shaping the transcriptional profiles of HSTCL cells, we performed RNA sequencing (RNA-seq) of the three murine and two human cell lines. Additionally, we integrated publicly available RNA-seq data from three primary HSTCL patient samples and one healthy control spleen sample.^24^ Differential gene expression analysis of the tumor cells compared to respective healthy controls, filtered for human-mouse homologs, revealed a considerable number of similarly dysregulated genes; a total of 255 overlapping differentially expressed genes were observed, including 191 genes co-upregulated and 64 genes co-downregulated across the mouse and human cell lines and the primary HSTCL patient tumors (**Figure 2A**). Gene set enrichment analysis (GSEA) revealed common significantly dysregulated pathways, with upregulated pathways including cell cycle, G2/M checkpoint and E2F targets (**Figure 2B,C**, **Supplemental Figure 4A**), indicating the highly proliferative nature of the tumor cells and reiterating STAT5B^N642H^ as a driver of cell-cycle progression. Notably, cell cycle was previously identified as the top enriched pathway of upregulated genes in HSTCL tumor cells compared with normal γδ T cells.^25^ In contrast, commonly downregulated pathways included TP53 targets and lymphocyte activation (**Figure 2B**). TCR pathway genes were significantly downregulated in both murine and human cell lines (**Figure 2C**, **Supplemental Figure 4B**), in line with the absence of surface CD3 and TCRδ expression on these cells (**Figure 1E**).

**Figure 2:**
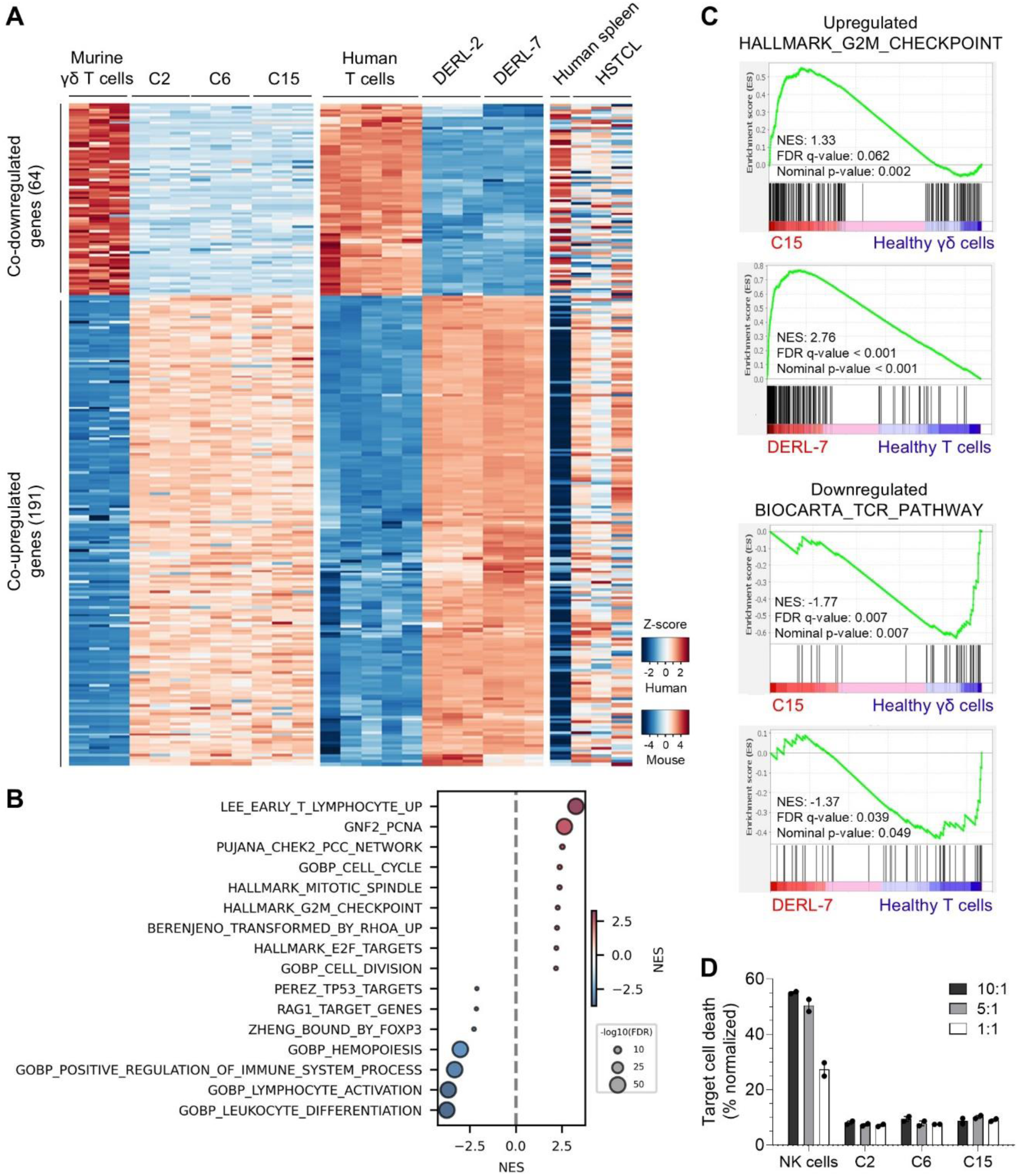
Murine γδ T-cell lymphoma cell lines have overlapping transcriptional profiles with human HSTCL cells and do not possess functional cytotoxic capacity. **A)** Heatmap of normalized expression (Z-scores) for commonly differentially expressed genes (DEGs) in murine C2, C6 and C15 cell lines (*n* = 3 each) compared to WT C57BL/6 γδ T cells (*n* = 3), human DERL-2 and DERL-7 cell lines (*n* = 3 each) compared to normal human T cells (*n* = 5), and primary human HSTCL patient samples (*n* = 3) compared to healthy spleen (*n* = 1), determined by RNA-seq (primary sample data were previously reported^24^). Genes were filtered for human–mouse homologs. **B)** Gene set enrichment analysis (GSEA) of overlapping DEGs across the murine and human cell lines and primary HSTCL patient samples, pre-ranked by log2 fold change, displayed as a bubble plot showing significantly up- (red) and down-regulated (blue) biological pathways; thresholds: |NES| > 2, FDR *q* < 0.05, nominal *p* < 0.01. Bubble size represents -log10 (FDR). **C)** GSEA of DEGs from the human DERL-7 and mouse C15 cell lines, displaying significantly up- and down-regulated biological pathways. NES, normalized enrichment score. **D)** Cell based killing assay examining the cytotoxic capacity of C2, C6 and C15 cells against the target cell line, YAC-1. Cells were seeded at 10:1, 5:1 and 1:1 ratios of γδTCL cells:target cells and incubated for 4 hr. Freshly isolated and IL-2 stimulated primary murine NK cells were used as a positive control. Target cell viability was measured by flow cytometry, and data are graphed as mean (± range) of technical duplicates, representative of three independent experiments (*n* = 3).

Given previous reports of negligible cytotoxic activity in the DERL-2 and DERL-7 HSTCL cell lines^18^ and of consistent downregulation of cytotoxic factors in primary HSTCL tumor cells,^25,26^ we determined the cytolytic potential of our murine γδ T-cell lines. Compared to the strong cytotoxic activity of primary murine NK cells against target YAC-1 tumor cells (up to 55% target cell killing), the C2, C6 and C15 cell lines displayed minimal cytotoxic capacity, inducing 5-10% target cell lysis irrespective of the effector:target cell ratio (**Figure 2D**).

Together, our findings demonstrate that STAT5B^N642H^-mediated transformation of murine γδ T cells can model various immunophenotypic, transcriptomic and functional features of HSTCL tumor cells.

### Murine C15 cell line allografts generate an aggressive HSTCL-like phenotype *in vivo*

We next investigated whether our murine cells could be used to generate an HSTCL-like disease phenotype *in vivo*. We transplanted C15 cells intravenously via the tail vein into immunodeficient NSG mice, which lack host T-, B- and NK cells. For specific tumor cell tracing, we analysed intracellular FLAG^+^ cells in the PB of recipient mice by flow cytometry over time, with tumor cells becoming detectable in PB at 5-7 weeks post-transplant (**Supplemental Figure 5A**), confirming PB involvement in this model. C15-recipient NSG mice succumbed to an aggressive γδTCL phenotype with 100% penetrance and a median survival of 70 days (**Figure 3A**). These mice developed hepatosplenomegaly characterized by significantly elevated liver and spleen weights (**Figure 3B**). Immunohistochemistry (IHC) revealed highly proliferative (Ki67^+^), CD3^+^ tumor cells infiltrating the liver (**Figure 3C**), as well as the spleen and BM (**Supplemental Figure 5B**). Importantly, flow cytometry of FLAG^+^ C15 tumor cells showed that CD3 and TCRδ cell surface expression were restored in the *in vivo* setting (**Supplemental Figure 5C**), in line with the immunophenotype of primary HSTCL tumors.^2^ To confirm that the allograft potential and *in vivo* disease phenotype of the C15 tumor cells are not an artefact of a single clone, we also allografted the polyclonal M2 cell line, from which the C15 clone was established, into NSG mice. M2-recipient mice rapidly developed a lethal γδTCL disease with 100% penetrance, a median survival of 31 days and evidence of hepatosplenomegaly (**Supplemental Figure 5D-F**).

**Figure 3:**
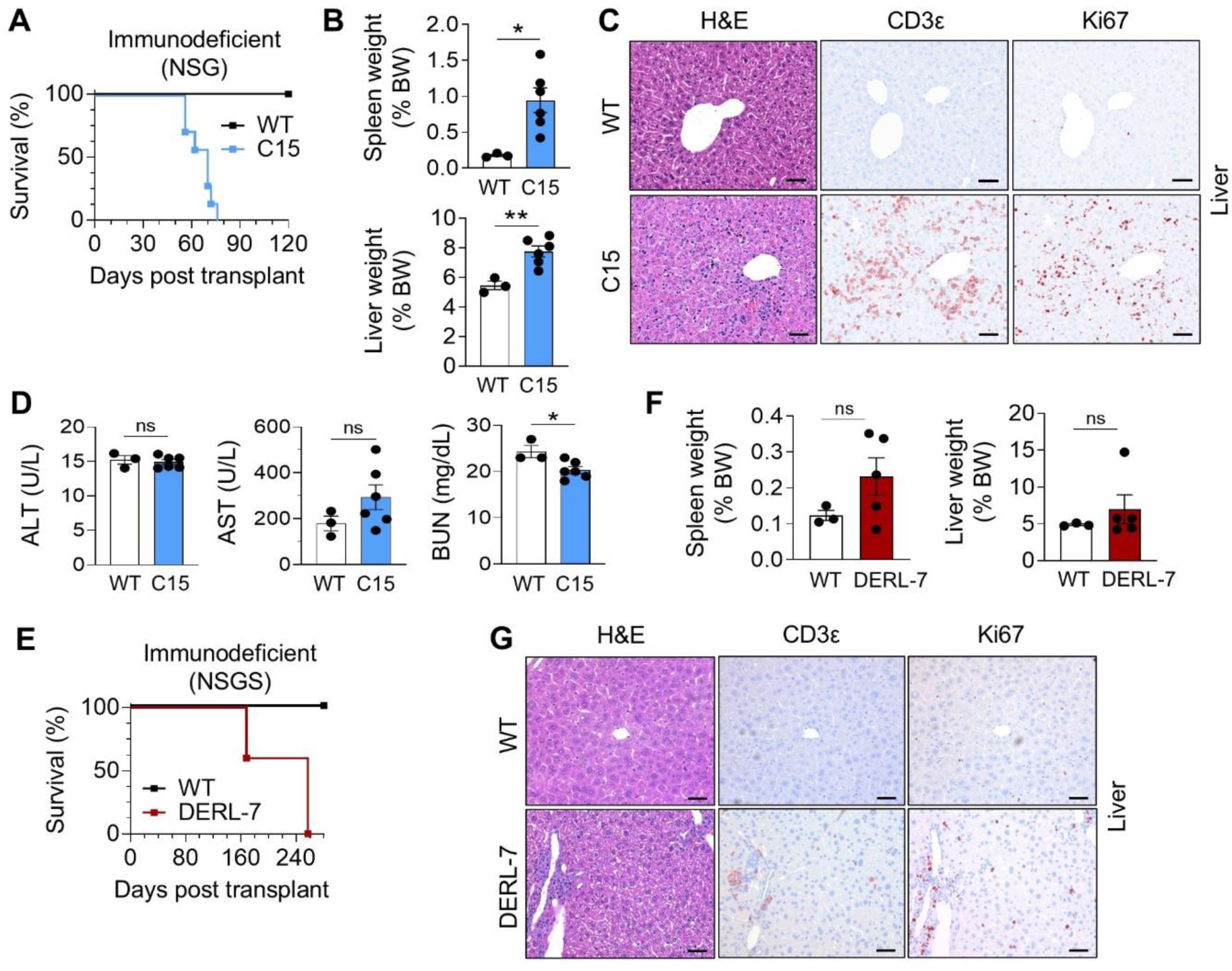
Murine C15 cell line intravenous allografts give rise to an HSTCL-like disease phenotype in immunodeficient mice. **A)** Kaplan-Meier survival analysis of NSG mice transplanted with C15 cells (*n* = 7) via tail vein injection, or control (WT) mice (*n* = 3). **B)** Spleen and liver weights (as % of body weight, BW) of WT and C15-recipient NSG mice. Data are graphed as mean (± SEM). \**p* < 0.05, \*\**p* < 0.01; unpaired two-tailed Student’s t-test. **C)** Representative images from immunohistochemistry (IHC) analysis of consecutive liver sections from WT or C15-recipient NSG mice, stained for CD3, Ki67 and H&E and imaged by light microscopy (scale bar = 50 μm). **D)** Levels of aspartate aminotransferase (AST), alanine aminotransferase (ALT) and blood urea nitrogen (BUN) in the plasma of WT (*n* = 3) and C15-recipient (*n* = 6) NSG mice. Data are graphed as mean (± SEM). \**p* < 0.05, ns = not significant; unpaired two-tailed Student’s t-test. **E)** Kaplan-Meier survival analysis of NSGS mice transplanted with DERL-7 cells (*n* = 5) via tail vein injection, or control (WT) mice (*n* = 3). **F)** Spleen and liver weights (as % of BW) of WT and DERL-7-recipient NSGS mice. Data are graphed as mean (± SEM). ns = not significant; unpaired two-tailed Student’s t-test. **G)** Representative images from IHC analysis of consecutive liver sections from WT or DERL-7-recipient NSGS mice, stained for CD3, Ki67 and H&E and imaged by light microscopy at 20x magnification (scale bar = 50 μm).

Further assessment of clinical features in the C15 model revealed a significant reduction in haemoglobin and platelet levels in the PB of C15-recipient mice compared with WT NSG mice (**Supplemental Figure 5G**), consistent with the frequent observations of anaemia and thrombocytopenia in HSTCL patients.^2^ No differences in body weight were observed in the tumor-burdened mice compared to control WT mice (**Supplemental Figure 5H**). We also observed indications of elevated ALT and AST liver transaminase levels in the plasma of C15-recipient mice, indicating liver damage (**Figure 3D**), as is also observed in HSTCL patients.^2^ Slightly reduced blood urea nitrogen (BUN) levels were found (**Figure 3D**), indicating normal kidney function and highlighting organ specificity of the malignant cells.

To the best of our knowledge, there have been no reports of systemic xenograft model establishment using human DERL cell lines. Therefore, for comparative purposes, we transplanted DERL-7 cells (*STAT5B^N642H^*homozygous) intravenously into immunodeficient NSGS mice, which express human IL3, GM-CSF (*CSF2*) and SCF (*KITLG*) to support human hematopoietic cell engraftment. All mice developed terminal disease, with a median survival of 257 days (**Figure 3E**). However, development of splenomegaly was inconsistent (3/5 recipient mice), and hepatomegaly was rare (1/5 recipient mice) (**Figure 3F**). IHC of the liver revealed minimal infiltration of Ki67^+^ CD3^+^ tumor cells (**Figure 3G**). Unlike allografted C15 cells, liver-infiltrating human CD45^+^ DERL-7 tumor cells remained negative for cell surface TCRδ and CD3 expression (**Supplemental Figure 6A**). Analysis of plasma ALT, AST and BUN levels in DERL-7-recipient mice revealed no significant differences compared to WT NSGS mice (**Supplemental Figure 6B**). Therefore, compared to human DERL-7 HSTCL xenografts, our murine C15 allografts have a more rapid disease onset and display more representative HSTCL disease phenotypes.

We next tested the allograft potential of C15 cells into immunocompetent mice, which would represent a major advantage of this model. We transplanted C15 cells (Ly5.2^+^) intravenously into immunocompetent (C57BL/6 Ly5.1) recipient mice, which became detectable in PB at 7 weeks post-transplant (**Supplemental Figure 7A**). C15-recipient Ly5.1 mice developed γδTCL disease with 80% disease penetrance and a median survival of 86 days (**Figure 4A**), displaying prominent hepatosplenomegaly (**Figure 4B**). Flow cytometry of liver cells confirmed a large expansion of Ly5.2^+^ γδ tumor cells (**Figure 4C**, **Supplemental Figure 7B**). As with the immunodeficient model, the C15 tumor cells re-expressed cell surface CD3 and TCRδ *in vivo* (**Supplemental Figure 7C**). Haemoglobin and platelet levels in the C15-recipient Ly5.1 mice were also reduced compared to WT Ly5.1 mice, and WBC numbers in the PB were elevated as expected given the leukemic involvement of the tumor cells (**Supplemental Figure 7D,E**). Elevated WBC counts in HSTCL patients are rare but were reported in patients coinciding with higher performance status and elevated LDH levels.^27^ C15-recipient Ly5.1 mice displayed a slight, albeit not significant, reduction in body weight gain compared to WT control mice (**Supplemental Figure 7F**). Comprehensive IHC analyses revealed that liver-infiltrating tumor cells were CD3^+^, TCRδ^+^, Ki67^+^, and could be specifically detected via staining for the FLAG-tag of the hSTAT5B^N642H^ transgenic protein (**Figure 4D**, **Supplemental Figure 7G**). Histological assessment of the spleen revealed diffuse and sinusoidal infiltration with disrupted architecture and considerable atrophy of the white pulp (**Supplemental Figure 7H,I**). A sinusoidal infiltration pattern was also found in the BM and liver with tumor cells also present in hepatic portal fields (**Figure 4D**, **Supplemental Figure 7I**), in line with observations in human HSTCL patients^1^. Across various organs, tumor cells were found infiltrating most prominently into the spleen, liver, BM, lymph nodes and lung, with minimal infiltration observed in the kidney, brain and heart (**Figure 4E, Supplemental Figure 7I**). In this immunocompetent model, we observed more highly elevated ALT and AST levels in the plasma of C15-recipient Ly5.1 mice, with no changes in BUN levels (**Figure 4F**).

**Figure 4:**
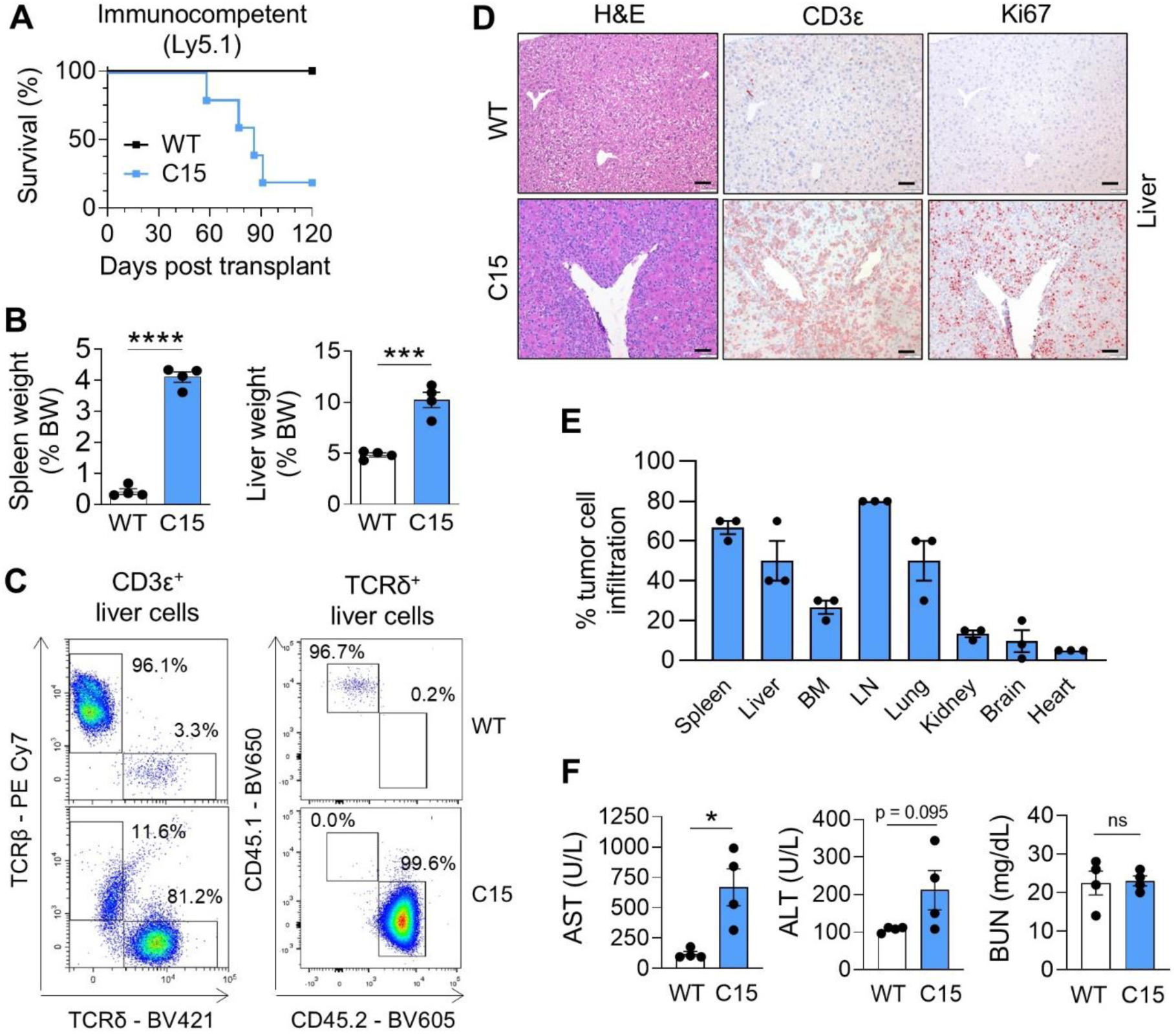
Murine C15 cell line intravenous allografts give rise to an HSTCL-like disease phenotype in immunocompetent mice. **A)** Kaplan-Meier survival analysis of C57BL/6 Ly5.1 mice transplanted with C15 cells (*n* = 5) via tail vein injection, or control (WT) mice (*n* = 4). **B)** Spleen and liver weights (as % of BW) of WT and diseased C15-recipient Ly5.1 mice. Data are graphed as mean (± SEM). \*\*\**p* < 0.001, \*\*\*\**p* < 0.0001; unpaired two-tailed Student’s t-test. **C)** Representative FACS plots showing percentages of TCRβ^+^ and TCRδ^+^ cells of CD3^+^ liver cells, and CD45.1^+^ and CD45.2^+^ cells of TCRδ^+^ liver cells, from WT and C15-recipient Ly5.1 mice. **D)** Representative images from IHC analysis of consecutive liver sections from WT or C15-recipient Ly5.1 mice, stained for CD3, Ki67 and H&E and imaged by light microscopy at 20x magnification (scale bar = 50 μm). **E)** Quantification of tumor cell infiltration (%) into various organs of C15-recipient Ly5.1 mice (*n* = 3), using representative IHC images of CD3 staining. **F)** Levels of AST, ALT and BUN in the plasma of WT and C15-recipient Ly5.1 mice. Data are graphed as mean (± SEM). \**p* < 0.05, ns = not significant; unpaired two-tailed Student’s t-test.

Taken together, we developed an HSTCL-like preclinical mouse model driven by oncogenic STAT5B^N642H^, which can be easily and rapidly generated using cell line allografts and that recapitulates key features of HSTCL in both immunocompetent and immunodeficient settings.

### JAK inhibitors display selective anti-tumor efficacy against murine and human HSTCL cell lines *in vitro*

We next utilized the HSTCL cell models to test potential targeted therapies for this aggressive disease. Whilst the STAT5B^N642H^ mutated protein is more persistently activated and more resistant to dephosphorylation than wildtype STAT5B,^14^ we and others have shown that it is not ‘constitutively’ active and that its activation is still reliant on upstream cytokine and JAK activity.^10,17^ In line, the murine and human HSTCL cell lines remain dependent on IL-2 signaling for proliferation and activation of the STAT5B oncogene (**Figure 1B,C, Supplemental Figure 2B-D**). Thus, we reasoned that blocking this upstream signaling via JAKi could serve as a promising targeted therapeutic strategy in *STAT5B*-mutated HSTCL, representing one-third of all HSTCL cases. Based on our RNA-seq data, both the murine and human HSTCL cell lines express transcripts for all four JAK kinases, with the murine tumor cells most highly expressing *Jak1* and *Jak3*, and the human cell lines most highly expressing *JAK1*. *Jak2*/*JAK2* is expressed to a very low level in all cell lines (**Supplemental Figure 8A**). We selected a panel of eight JAKi with various JAK selectivity profiles (**Figure 5A**), many of which are clinically approved, and used these compounds for *in vitro* testing on the murine and human HSTCL cell lines in the presence of IL-2. The JAKi were effective in reducing the viability of HSTCL cells after 48 hr, showing strong selectivity over two other human T-cell lymphoma cell lines, HH and Karpas 384, that do not harbor activating *JAK-STAT* mutations^19^ nor display detectable STAT5 activity (**Figure 5A**, **Supplemental Figure 8B,C**). As a control, we observed weak effects of a PARP inhibitor, olaparib, on reducing HSTCL cell viability by 48 hr (**Figure 5A**, **Supplemental Figure 8B**), indicating that *STAT5B*-mutant HSTCL cells are selectively sensitive to JAKi likely due to their dependence on downstream STAT5B signaling. The most effective JAKi, upadacitinib, strongly and selectively reduced the viability of all murine and human HSTCL cell lines with IC_50_ values between 27 and 102 nM (**Figure 5A,B**). This was paralleled by a dose-dependent reduction in STAT5 activation already after 4 hr of upadacitinib treatment as low as 10 nM, with an almost complete absence of STAT5 activity in all cell lines at 5 μM (**Figure 5C,D**, **Supplemental Figure 8D,E**). We confirmed this effect using a second JAKi, tofacitinib (cell viability IC_50_ values between 443 and 645 nM), observing diminished STAT5 activity in C15 cells after 4 hr of treatment (**Figure 5E**), validating the targeted approach of using JAKi to block downstream activation of the STAT5B oncogene.

**Figure 5:**
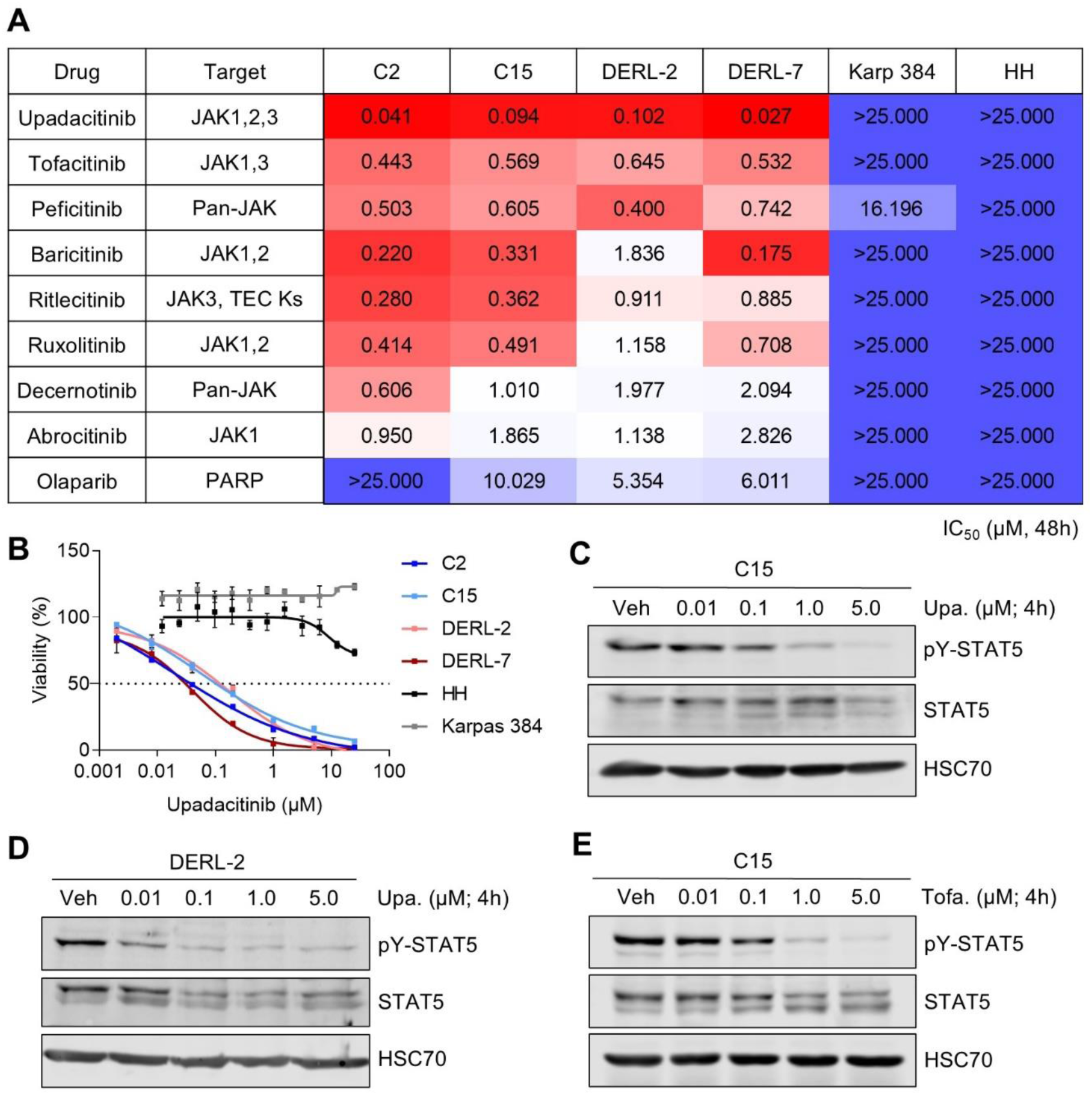
HSTCL cell lines are selectively sensitive to JAK inhibitors *in vitro*. **A)** Heatmap of IC_50_ values calculated from drug-response analyses using CellTiter-Blue viability assays upon 48 hr treatment of the indicated drugs in IL-2 supplemented media. Average IC_50_ values from three independent experiments are shown (*n* = 3). TEC Ks: Tec kinases. **B)** Cell viability curves upon 48 hr upadacitinib treatment at various concentrations using murine (C2, C15) and human (DERL-2, DERL-7) HSTCL cell lines, and control human (HH, Karpas 384) TCL cell lines. Data are graphed as mean (± SD) of technical triplicates from one experiment, representative of three independent experiments (*n* = 3). **C-E)** Western blots showing STAT5 activity in C15 cells (C) or DERL-2 cells (D) treated with upadacitinib, or C15 cells treated with tofacitinib (E), for 4 hr at the indicated concentrations. During this time, cells were starved of IL-2 for 3.5 hr and then restimulated with 10 ng/mL IL-2 for 30 min. Immunoblotting for pY-STAT5 and total STAT5 was performed, with HSC70 serving as a loading control (blots representative of two to three independent experiments each; *n* = 2-3). Veh, vehicle (DMSO).

### The JAK inhibitor upadacitinib reduces oncogenic STAT5 activity and HSTCL disease burden *in vivo*

With these promising *in vitro* results, we selected the top performing JAK1/2/3 inhibitor upadacitinib for further *in vivo* testing using our HSTCL preclinical model. NSG mice were transplanted intravenously with C15 cells and after 10 days, to allow tumor cell engraftment, mice were treated by oral gavage once daily (five days per week) with either upadacitinib (10 mg/kg) or vehicle. At eight weeks post-transplant, a vehicle-treated mouse displayed symptoms of terminal disease, at which point all mice were sacrificed and a comparative endpoint analysis was performed. Notably, we observed reduced liver and spleen weights in upadacitinib treated mice (**Figure 6A**, **Supplemental Figure 9A**), and reduced tumor burden (FLAG^+^ cells) in the liver, BM, spleen and PB as determined by flow cytometry, compared to vehicle treated mice (**Figure 6B**, **Supplemental Figure 9B**). Reduced tumor cell burden was also visualized in liver and spleen tissues by IHC (**Figure 6C**). The drug was well tolerated, and we did not observe any impact of toxicity on haemoglobin or platelet levels in treated mice (**Supplemental Figure 9C**). We assessed STAT5 activation levels in splenocytes and observed a strong reduction in activated STAT5 in upadacitinib treated mice compared to vehicle treated mice (**Figure 6D**). Together, these data confirm that the STAT5B oncogene-driven tumor cells remain dependent on upstream JAK signaling in an *in vivo* setting, supporting the use of upadacitinib as a targeted therapy for *STAT5B*-mutant HSTCL.

**Figure 6:**
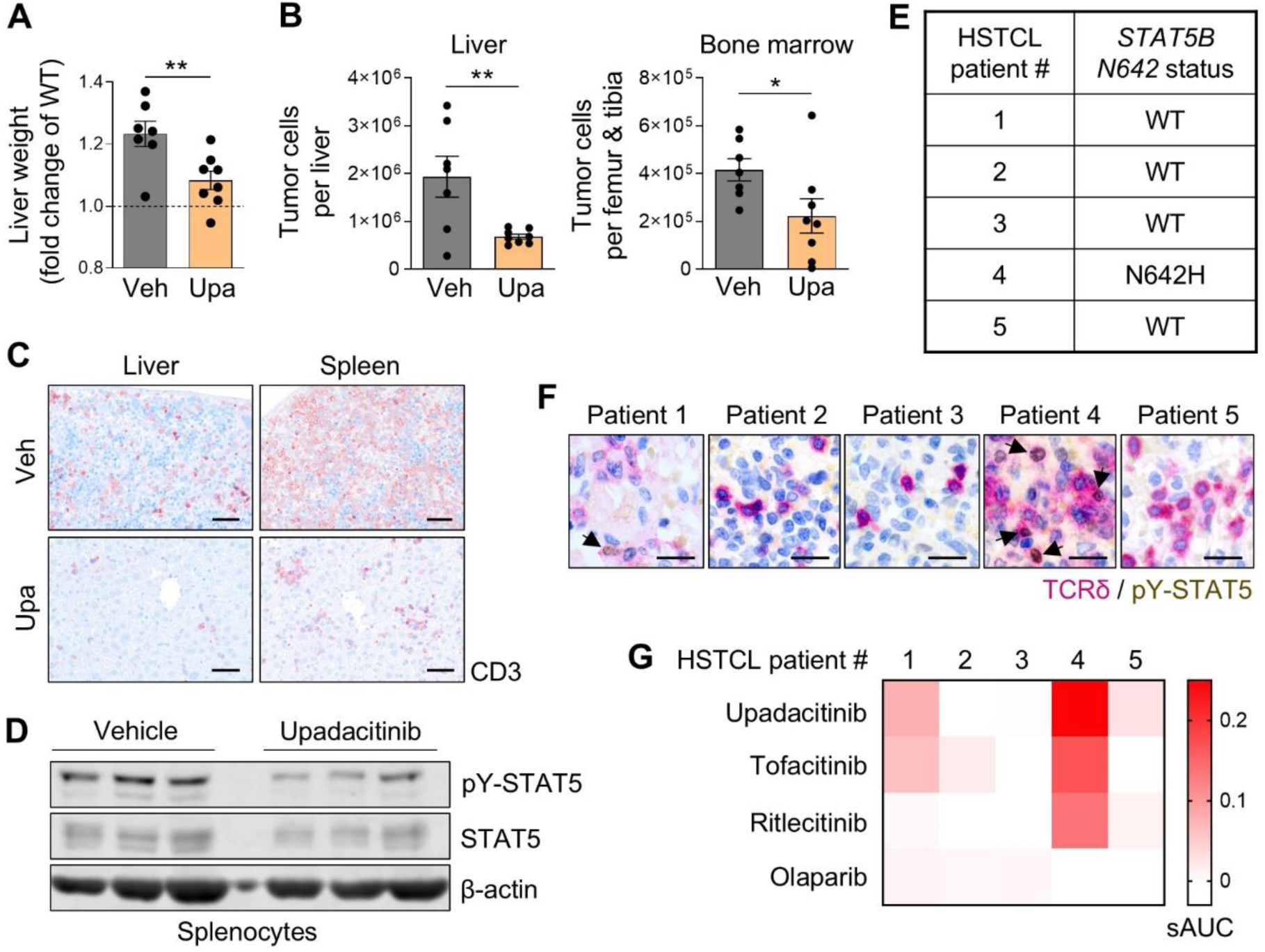
JAK1/2/3 inhibitor, upadacitinib, displays anti-tumor efficacy in the C15 allograft *in vivo* model and in primary HSTCL patient cells harboring the STAT5B^N642H^ mutation. **A)** Liver weights of NSG mice transplanted with C15 cells and treated with vehicle (veh; *n* = 7) or upadacitinib (*n* = 8), graphed as fold change of average liver weight of WT NSG mice. All mice were analysed 56 days post-transplant. Data are graphed as mean (± SEM). \*\**p* < 0.01; unpaired two-tailed Student’s t-test. **B)** Absolute tumor cell numbers (intracellular FLAG+ cells) in the liver and bone marrow of vehicle or upadacitinib treated C15-recipient NSG mice, analysed by flow cytometry. Data are graphed as mean (± SEM). \**p* < 0.05, \*\**p* < 0.01; unpaired two-tailed Student’s t-test. **C)** Representative images from IHC analysis of liver and spleen sections from vehicle or upadacitinib treated C15-recipient NSG mice, stained for CD3 and imaged by light microscopy at 20x magnification (scale bar = 50 μm). **D)** Western blot showing STAT5 activity in splenocytes from vehicle or upadacitinib treated C15-recipient NSG mice (*n* = 3; three individual mice are shown per treatment group, representative of all mice from each group). Immunoblotting for pY-STAT5 and total STAT5 was performed, with β-actin serving as a loading control. **E)** Table indicating the *STAT5B* mutation status at position N642 of HSTCL patients (*n* = 5), determined by digital PCR. **F)** Representative images from IHC analysis of TCRδ (magenta) and pY-STAT5 (brown) double staining in the bone marrow (patient 1) or spleen (patients 2-5) of HSTCL patients, imaged by light microscopy (scale bar = 20 μm). Black arrows indicate tumor cells with TCRδ and pY-STAT5 double staining. **G)** Heatmap of selective area under the curve (sAUC) values calculated from drug-response viability analyses using flow cytometry upon 24 hr treatment of primary HSTCL patient cells with the indicated drugs (concentration range: 0.001-10 µM). Average AUC values of normal and tumor cell populations from technical duplicate samples were used.

### Primary HSTCL patient tumor cells with a *STAT5B^N642H^* mutation display STAT5 activity and sensitivity to JAK inhibition *ex vivo*

Next, we assessed the sensitivity of primary HSTCL patient samples to JAKi. Given the rarity of the disease, we managed to obtain both viable and FFPE tumor samples from five HSTCL patients (**Supplemental Table 1**). To determine the relevance of *STAT5B* mutation status to JAKi sensitivity, we screened the patient material for the presence of the *STAT5B^N642H^* mutation by digital PCR using a mutant allele-specific probe. The high sensitivity and selectivity of the assay to the *STAT5B^N642H^* mutation was confirmed using DERL-2, DERL-7 and Karpas 384 cell lines (**Supplemental Figure 10A-C**). Using the assay, we detected the *STAT5B^N642H^* mutation in one of the five HSTCL patients (20%; **Figure 6E, Supplemental Figure 10D**). To further assess whether *STAT5B^N642H^* mutation status correlates with STAT5 activation in HSTCL tumor cells, we performed pY-STAT5/TCRδ double staining by IHC using FFPE tumor sections. Patient 4, harboring the *STAT5B^N642H^*mutation, displayed clear nuclear STAT5 activity across TCRδ^+^ tumor cells. Weak STAT5 activity was detected in some tumor cells from patient 1, but was absent in tumor cells from patients 2, 3 and 5 (**Figure 6F, Supplemental Figure 11**).

We then incubated viable HSTCL patient samples with varying concentrations of upadacitinib, tofacitinib or ritlecitinib *ex vivo* for 24 hr, and measured cell viability using flow cytometry. The patient samples included a mixture of both tumor cells and non-malignant microenvironmental cells, which could be distinguished using defined surface markers (**Supplementary Table 1**). This allowed us to determine the effects of the JAKi on both tumor and healthy cell populations, to generate selective area under the curve (sAUC) values (tumor cell AUC minus healthy cell AUC) as an indication of the selective anti-tumor effect of the drugs. Of note, a 24-hr incubation time was selected for the primary samples, as opposed to the 48-hr JAKi treatment of the cell lines *in vitro*, due to a considerable loss of cell viability of vehicle-treated patient cells beyond 24 hr of *ex vivo* culture. Strikingly, upon JAKi treatment we observed a selective reduction in viability of the *STAT5B^N642H^*-positive tumor cells (patient 4), with the strongest sensitivity observed for upadacitinib (**Figure 6G**), which induced cell death in >40% of tumor cells by 24 hr at 100 nM (**Supplemental Figure 12A**). Interestingly, patient 1, who displayed low-level STAT5 activity in some tumor cells, showed weak sensitivity towards JAKi treatment, albeit less than patient 4. There was no impact on the viability of tumor cells from the other *STAT5B* WT patients or on healthy cells from all patients (**Figure 6G, Supplemental Figure 12A-C**). As a control, the PARP inhibitor, olaparib, did not impact the viability of any patient cell population (**Figure 6G, Supplemental Figure 12D**).

Overall, these data suggest that JAKi, such as upadacitinib, could represent a beneficial, targeted treatment for HSTCL patients harboring activating *STAT5B* mutations.

## Discussion

Recent reports on the incidence of HSTCL indicate that cases are rising whilst, alarmingly, the overall survival has not significantly improved over the last decades.^28^ There is an urgent need for faithful preclinical models and the testing of new therapeutic strategies for this orphan disease. Here, we have developed a STAT5B-driven mouse model, easily and robustly derived using cell line allografts, which recapitulates various clinical, immunophenotypic and genomic features of HSTCL. We have utilized this *in vivo* model, together with patient derived HSTCL cell lines and primary samples, to demonstrate the selective anti-tumor efficacy of the JAKi upadacitinib against HSTCL cells with an activating *STAT5B* mutation, representing one-third of HSTCL patients. Our findings could have immediate clinical implications given that JAKi, such as upadacitinib, are already approved for the treatment of various indications (e.g. rheumatoid arthritis, atopic dermatitis and ulcerative colitis) and could be rapidly repurposed for clinical testing in HSTCL patients. While the number of HSTCL patient samples used in this study is relatively low, in line with the rarity of the disease, our findings indicate that *STAT5B* mutation status or the status of STAT5 protein activity could potentially serve as markers for predicting JAKi treatment response, warranting further preclinical and clinical validation.

With the high frequency of JAK-STAT pathway hyperactivation in HSTCL and across most T-cell lymphoma entities,^6,29^ interest in targeted therapies to block this pathway is increasing.^30,31^ However, the use of JAKi, particularly upadacitinib, in treating HSTCL patients currently remains unexplored. Moskowitz *et al.* reported on a phase II biomarker-driven clinical trial assessing the efficacy of JAK1/2 inhibitor ruxolitinib in patients with various peripheral T-cell leukemias/lymphomas (PTCLs) harboring JAK-STAT mutations.^32^ Overall, a clinical benefit was achieved in 53% of the ruxolitinib treated PTCL patients. Two HSTCL patients were included in this study, both with *STAT5B^N642H^* mutations, and one patient achieved a partial response.^32^ A more recent phase I/II study of the selective JAK1 inhibitor, golidocitinib, in relapsed or refractory PTCL revealed an overall objective response in 44.3% of patients.^33^ However, no HSTCL patients were included in this study. Case reports have indicated the efficacy of upadacitinib treatment in inducing clinical responses in patients with cutaneous T-cell lymphoma (CTCL).^34,35^ These studies demonstrate the potential of JAKi as targeted therapies for PTCL patients, particularly those with hyperactivating *JAK-STAT* mutations, and, together with our findings, warrant the clinical testing of upadacitinib in HSTCL patients. Interestingly, while we observed a strongly selective response to JAKi from the HSTCL patient with *STAT5B^N642H^*, we also observed some JAKi sensitivity from the tumor cells of patient 1, who was negative for this mutation but still displayed some nuclear STAT5 activity. Therefore, screening for STAT5 activation status in tumor cells by IHC may represent a beneficial functional marker in addition to *STAT5B* mutation status to predict patient response to JAKi. Future preclinical experiments testing upadacitinib in combination with commonly used chemotherapeutic agents (e.g. components of ICE-based regimens) would also be valuable to explore synergistic effects and further improved efficacy.

In our *in vitro* cell viability assays, upadacitinib displayed the strongest anti-tumor activity across all HSTCL tumor cell lines tested, with the lowest IC_50_ values out of the eight JAKi screened. For this reason, we chose upadacitinib for further preclinical testing *in vivo*. Upadacitinib (Rinvoq; ABT-494) is widely reported as a JAK1-selective inhibitor.^36–38^ However, detailed studies comparing JAK family target specificity of different JAKi in human PBMCs and whole blood revealed that, while the greatest potency was observed against JAK1, inhibition of JAK2- and JAK3-dependent signaling by upadacitinib was observed at clinically relevant doses.^39,40^ From our JAKi screening results on HSTCL cell viability, it therefore appears that, at least *in vitro*, the most effective JAKi were those with broader JAK selectivity profiles or that include inhibition of both JAK1 and JAK3 signaling. This is perhaps expected given the important role of JAK1/3 in mediating lymphocyte proliferation and homeostasis downstream of IL-2 and other cytokines acting via the common γ-chain receptor.^41^ Further support that the efficacy of upadacitinib in reducing HSTCL cell viability may result from its inhibition of both JAK1 and JAK3 comes from the relatively high IC_50_ values of the JAK1-specific inhibitor, abrocitinib (IC_50_ values: abrocitinib, 0.95 - 2.83 μM; upadacitinib, 0.03 - 0.10 μM). Abrocitinib has a >340-fold higher affinity for JAK1 over JAK3, with an IC_50_ for JAK1 of 29 nM.^42^ Further genetic and pharmacologic approaches are required to dissect the individual involvement of JAK1 and JAK3 in HSTCL pathogenesis.

To the best of our knowledge, we describe here the first preclinical mouse model recapitulating an HSTCL-like disease phenotype, which can be modelled in both an immunodeficient and immunocompetent setting. Li *et al.* previously reported that transgenic mice deficient in the inhibitory helix-loop-helix protein Id3 (Id3^-/-^) spontaneously develop a γδTCL disease with a penetrance of 27%.^43^ Adoptive transfer of the tumor cells into immunodeficient Rag1^-/-^ recipients resulted in a fully-penetrant disease with broad organ infiltration and a median survival of around 16 weeks. While the authors state that this model may represent a tool to study HSTCL disease, the similarities of the model to human HSTCL were not extensively assessed. Disease development in immunocompetent mice was also not assessed in this study.^43^

Activating *STAT5B* mutations, particularly *STAT5B^N642H^*, are also found in other γδTCL entities, including primary cutaneous γδ T-cell lymphoma (PCGDTL), TCRγδ^+^ monomorphic epitheliotropic intestinal T-cell lymphoma (MEITL) and γδ T cell large granular lymphocytic leukemia (T-LGLL).^6^ While the murine tumor cells could also serve as tools to study oncogenic STAT5B in these diseases, our profiling indicates that the models share a high degree of similarity to HSTCL and are more divergent from these other entities. For example, we did not observe strong infiltration of the C15 tumor cells into the skin or intestine, which are the main tumor sites for PCGDTL and MEITL, respectively. Additionally, splenomegaly and hepatomegaly are infrequent in γδ T-LGLL cases, and the disease course is relatively indolent.^6,44,45^

While we have demonstrated that our mouse cell lines possess various features in common with human HSTCL cells, even human HSTCL cell lines *in vitro* display considerable differences to primary HSTCL tumor cells. Notably, the downregulation of surface CD3 and TCRδ is highly uncharacteristic of the primary disease,^2^ emphasizing the limitations of 2D cell culture models and the importance of faithful *in vivo* models. Importantly, we observed dynamic restoration of CD3 and TCRδ expression on the surface of the murine C15 tumor cells when profiled *ex vivo*. It is unclear whether this dynamic regulation is mediated by the STAT5B^N642H^ oncogene, although we have recently demonstrated in a T-ALL mouse model driven by this mutant, that STAT5B^N642H^ promotes downstream TCR signaling even in the absence of surface TCR.^16^ This mimicking of TCR signaling may render surface TCR and co-receptor components redundant and induce a negative feedback signal to downregulate them, particularly *in vitro* where antigenic stimulus is absent and STAT5-activating IL-2 levels are relatively high.

Overall, STAT5B^N642H^-induced transformation of murine γδ T cells and *in vivo* modelling can recapitulate various phenotypic and genomic features of human HSTCL. This model serves as a valuable tool for mechanistic studies and urgently needed preclinical testing of new therapeutic strategies, where we reveal here the selective anti-tumor efficacy of the JAKi upadacitinib in HSTCL with hyperactive STAT5B. Preclinical studies using this model could also inform on new treatment options relevant for other aggressive γδTCL entities where oncogenic *STAT5B* mutations are prevalent.

## References

1. Yabe M, Miranda RN, Medeiros LJ. Hepatosplenic T-cell Lymphoma: a review of clinicopathologic features, pathogenesis, and prognostic factors. Hum Pathol. 2018;74:5–16.

2. Pro B, Allen P, Behdad A. Hepatosplenic T-cell lymphoma: a rare but challenging entity. Blood. 2020;136(18):2018–2026.

3. Stuver R, Epstein-Peterson ZD, Horwitz SM. Few and far between: clinical management of rare extranodal subtypes of mature T-cell and NK-cell lymphomas. Haematologica. 2023;108(12):3244–3260.

4. Armitage JO. The aggressive peripheral T-cell lymphomas: 2017. Am J Hematol. 2017;92(7):706–715.

5. Foss FM, Horwitz SM, Civallero M, et al. Incidence and outcomes of rare T cell lymphomas from the T Cell Project: hepatosplenic, enteropathy associated and peripheral gamma delta T cell lymphomas. Am J Hematol. 2020;95(2):151–155.

6. Schönefeldt S, Wais T, Herling M, et al. The Diverse Roles of gammadelta T Cells in Cancer: From Rapid Immunity to Aggressive Lymphoma. Cancers (Basel). 2021;13(24):6212.

7. Alonsozana EL, Stamberg J, Kumar D, et al. Isochromosome 7q: the primary cytogenetic abnormality in hepatosplenic gammadelta T cell lymphoma. Leukemia. 1997;11(8):1367–1372.

8. Lewis NE, Zhou T, Dogan A. Biology and genetics of extranodal mature T-cell and NKcell lymphomas and lymphoproliferative disorders. Haematologica. 2023;108(12):3261–3277.

9. McKinney M, Moffitt AB, Gaulard P, et al. The Genetic Basis of Hepatosplenic T-cell Lymphoma. Cancer Discov. 2017;7(4):369–379.

10. Kucuk C, Jiang B, Hu X, et al. Activating mutations of STAT5B and STAT3 in lymphomas derived from gammadelta-T or NK cells. Nat Commun. 2015;6:6025.

11. Nicolae A, Xi L, Pittaluga S, et al. Frequent STAT5B mutations in gammadelta hepatosplenic T-cell lymphomas. Leukemia. 2014;28(11):2244–2248.

12. Rajala HL, Eldfors S, Kuusanmaki H, et al. Discovery of somatic STAT5b mutations in large granular lymphocytic leukemia. Blood. 2013;121(22):4541–4550.

13. Bandapalli OR, Schuessele S, Kunz JB, et al. The activating STAT5B N642H mutation is a common abnormality in pediatric T-cell acute lymphoblastic leukemia and confers a higher risk of relapse. Haematologica. 2014;99(10):E188–E192.

14. de Araujo ED, Erdogan F, Neubauer HA, et al. Structural and functional consequences of the STAT5B(N642H) driver mutation. Nat Commun. 2019;10(1):2517.

15. Potdar S, Ianevski F, Ianevski A, et al. Breeze 2.0: an interactive web-tool for visual analysis and comparison of drug response data. Nucleic Acids Res. 2023;51(W1):W57–W61.

16. Suske T, Sorger H, Manhart G, et al. Hyperactive STAT5 hijacks T cell receptor signaling and drives immature T cell acute lymphoblastic leukemia. J Clin Invest. 2024;134(8):e168536.

17. Pham HTT, Maurer B, Prchal-Murphy M, et al. STAT5BN642H is a driver mutation for T cell neoplasia. J Clin Invest. 2018;128(1):387–401.

18. Di Noto R, Pane F, Camera A, et al. Characterization of two novel cell lines, DERL-2 (CD56+/CD3+/Tcry5+) and DERL-7 (CD56+/CD3-/TCRgammadelta-), derived from a single patient with CD56+ non-Hodgkin’s lymphoma. Leukemia. 2001;15(10):1641–1649.

19. Ng SY, Yoshida N, Christie AL, et al. Targetable vulnerabilities in T- and NK-cell lymphomas identified through preclinical models. Nat Commun. 2018;9(1):2024.

20. Dufva O, Kankainen M, Kelkka T, et al. Aggressive natural killer-cell leukemia mutational landscape and drug profiling highlight JAK-STAT signaling as therapeutic target. Nat Commun. 2018;9(1):1567.

21. John S, Robbins CM, Leonard WJ. An IL-2 response element in the human IL-2 receptor alpha chain promoter is a composite element that binds Stat5, Elf-1, HMG-I(Y) and a GATA family protein. EMBO J. 1996;15(20):5627–5635.

22. Li P, Mitra S, Spolski R, et al. STAT5-mediated chromatin interactions in superenhancers activate IL-2 highly inducible genes: Functional dissection of the Il2ra gene locus. Proc Natl Acad Sci U S A. 2017;114(46):12111–12119.

23. Heilig JS, Tonegawa S. Diversity of murine gamma genes and expression in fetal and adult T lymphocytes. Nature. 1986;322(6082):836–840.

24. Finalet Ferreiro J, Rouhigharabaei L, Urbankova H, et al. Integrative genomic and transcriptomic analysis identified candidate genes implicated in the pathogenesis of hepatosplenic T-cell lymphoma. PLoS One. 2014;9(7):e102977.

25. Song W, Zhang H, Yang F, et al. Single cell profiling of gammadelta hepatosplenic T-cell lymphoma unravels tumor cell heterogeneity associated with disease progression. Cell Oncol (Dordr). 2023;46(1):211–226.

26. Travert M, Huang Y, de Leval L, et al. Molecular features of hepatosplenic T-cell lymphoma unravels potential novel therapeutic targets. Blood. 2012;119(24):5795–5806.

27. Belhadj K, Reyes F, Farcet JP, et al. Hepatosplenic γδ T-cell lymphoma is a rare clinicopathologic entity with poor outcome:: report on a series of 21 patients. Blood. 2003;102(13):4261–4269.

28. Durani U, Go RS. Incidence, clinical findings, and survival of hepatosplenic T-cell lymphoma in the United States. Am J Hematol. 2017;92(6):E99–E101.

29. Fiore D, Cappelli LV, Broccoli A, Zinzani PL, Chan WC, Inghirami G. Peripheral T cell lymphomas: from the bench to the clinic. Nat Rev Cancer. 2020;20(6):323–342.

30. Waldmann TA. JAK/STAT pathway directed therapy of T-cell leukemia/lymphoma: Inspired by functional and structural genomics. Mol Cell Endocrinol. 2017;451(C):66–70.

31. Vahabi SM, Bahramian S, Esmaeili F, et al. JAK Inhibitors in Cutaneous T-Cell Lymphoma: Friend or Foe? A Systematic Review of the Published Literature. Cancers. 2024;16(5):861.

32. Moskowitz AJ, Ghione P, Jacobsen E, et al. A phase 2 biomarker-driven study of ruxolitinib demonstrates effectiveness of JAK/STAT targeting in T-cell lymphomas. Blood. 2021;138(26):2828–2837.

33. Song Y, Malpica L, Cai Q, et al. Golidocitinib, a selective JAK1 tyrosine-kinase inhibitor, in patients with refractory or relapsed peripheral T-cell lymphoma (JACKPOT8 Part B): a single-arm, multinational, phase 2 study. Lancet Oncol. 2024;25(1):117–125.

34. Kook H, Park SY, Hong N, et al. Severely pruritic mycosis fungoides successfully treated with upadacitinib. J Dtsch Dermatol Ges. 2024;22(3):450–451.

35. Castillo DE, Romanelli P, Lev-Tov H, Kerdel F. A case of erythrodermic mycosis fungoides responding to upadacitinib. JAAD Case Rep. 2022;30:91–93.

36. Parmentier JM, Voss J, Graff C, et al. In vitro and in vivo characterization of the JAK1 selectivity of upadacitinib (ABT-494). BMC Rheumatol. 2018;2:23.

37. Tanaka Y, Luo YM, O’Shea JJ, Nakayamada S. Janus kinase-targeting therapies in rheumatology: a mechanisms-based approach. Nat Rev Rheumatol. 2022;18(3):133–145.

38. Liu C, Kieltyka J, Fleischmann R, Gadina M, O’Shea JJ. A Decade of JAK Inhibitors: What Have We Learned and What May Be the Future? Arthritis Rheumatol. 2021;73(12):2166–2178.

39. Traves PG, Murray B, Campigotto F, Galien R, Meng A, Di Paolo JA. JAK selectivity and the implications for clinical inhibition of pharmacodynamic cytokine signalling by filgotinib, upadacitinib, tofacitinib and baricitinib. Ann Rheum Dis. 2021;80(7):865–875.

40. McInnes IB, Byers NL, Higgs RE, et al. Comparison of baricitinib, upadacitinib, and tofacitinib mediated regulation of cytokine signaling in human leukocyte subpopulations. Arthritis Res Ther. 2019;21(1):183.

41. Rochman Y, Spolski R, Leonard WJ. New insights into the regulation of T cells by γc family cytokines. Nat Rev Immunol. 2009;9(7):480–490.

42. Vazquez ML, Kaila N, Strohbach JW, et al. Identification of N-{cis-3-[Methyl(7H-pyrrolo[2,3-d]pyrimidin-4-yl)amino]cyclobutyl}propane-1-sulfonamide (PF-04965842): A Selective JAK1 Clinical Candidate for the Treatment of Autoimmune Diseases. J Med Chem. 2018;61(3):1130–1152.

43. Li J, Maruyama T, Zhang P, et al. Mutation of inhibitory helix-loop-helix protein Id3 causes γδ T-cell lymphoma in mice. Blood. 2010;116(25):5615–5621.

44. Sandberg Y, Almeida J, Gonzalez M, et al. TCRgammadelta+ large granular lymphocyte leukemias reflect the spectrum of normal antigen-selected TCRgammadelta+ T-cells. Leukemia. 2006;20(3):505–513.

45. Barila G, Grassi A, Cheon H, et al. Tgammadelta LGLL identifies a subset with more symptomatic disease: analysis of an international cohort of 137 patients. Blood. 2023;141(9):1036–1046.

