## Supplementary file for "A STAT5B-driven mouse model of hepatosplenic γδ T-cell lymphoma reveals therapeutic efficacy of JAK inhibition"

This file includes Supplementary Methods, Supplementary Tables 1-6, Supplementary Figures 1-12, Supplementary Figure Legends and Supplementary References.

### Supplementary Methods

#### hSTAT5B$^{N642H}$ transgenic mouse model

The *Vav1*-hSTAT5B$^{N642H}$ transgenic mouse model, from which the polyclonal and clonal γδ T-cell lines were derived, was previously generated as described elsewhere.[1] Briefly, the *Vav1*-hematopoietic vector containing human STAT5B$^{N642H}$ with a C-terminal FLAG-tag was digested with the *HindIII* restriction enzyme and gel purified for pronuclear injection into C57BL/6NCrl mice. The transgenic mice were identified by genotyping PCR and confirmed to express low-level (2x endogenous STAT5 levels), strongly activated STAT5B protein by Western blot of lymph node and spleen. Given the rapid development of CD8$^+$ T-cell disease in the hSTAT5B$^{N642H}$ transgenic mice, the colony must be propagated via *in vitro* fertilization with archived sperm cells.

#### Cell culture

Murine γδ T-cell lines were cultured in media containing RPMI 1640 supplemented with 20% hi-FBS (Biowest), 2 mM L-glutamine (Gibco), 10 U/ml penicillin/streptomycin (Biowest), 1X MEM Non-Essential Amino Acids Solution (Gibco), 20 mM HEPES buffer (Gibco), 1 mM sodium pyruvate (Gibco), 55 μM β-mercaptoethanol (Gibco) and 10 ng/ml IL-2 (ImmunoTools GmbH). The cell lines were sub-cultured every 2-3 days in fresh media using a standard trypsinization protocol. DERL-2 and DERL-7 human HSTCL cell lines and the HH cutaneous T-cell lymphoma cell line were obtained from the German Collection of Microorganisms and Cell Cultures GmbH (DSMZ). The Karpas 384 primary cutaneous γδ T-cell lymphoma cell line was obtained from the European Collection of Cell Cultures (ECACC). All human cell lines were cultured in media containing RPMI 1640 supplemented with 10% hi-FBS, 10 U/ml penicillin/streptomycin and 2 mM L-glutamine. DERL-2 and DERL-7 culture media was additionally supplemented with 10 ng/ml IL-2 (ImmunoTools GmbH). The murine YAC-1 lymphoma cell line was purchased from the American Type Culture Collection (ATCC) and was cultured in RPMI 1640 media including 10% FBS, 10 U/ml penicillin/streptomycin and 55 µM β-mercaptoethanol. Primary mouse NK cells were expanded in RPMI 1640 media containing 10% FBS, 10 U/ml penicillin/streptomycin, 55 µM β-mercaptoethanol and 3000 U/ml recombinant human IL2 (Proleukin, Sterimax).

All cell lines were incubated at 37°C with 5% $CO_2$ in a humidified environment and were regularly tested and confirmed negative for mycoplasma using PhoenixDx Mycoplasma Mix (Procomcure Biotech). The authenticity of the DERL-2, DERL-7, HH and Karpas 384 cell lines was confirmed by short tandem repeat (STR) profiling (Microsynth GmbH). Upon thawing cryopreserved cell lines, new stocks were prepared and frozen within 3-5 passages. Cell lines were kept in culture for no more than two months after resuscitation.

For cytokine starvation experiments, murine γδ T-cell lines were cultured for 48 hr in standard media with 10 ng/ml IL-2 (pre-starvation). Cells were then washed three times with Dulbecco's Phosphate-Buffered Saline (DPBS; Sigma-Aldrich) and cultured for an additional 8 hr in media without IL-2 (starvation). Cells were re-stimulated for 1 hr with 10 ng/ml IL-2 (re-stimulation). From all conditions, a cell pellet was collected for Western blot analysis.

**Western blotting**

For immunoblotting of cytokine-dependent cell lines, cells were stimulated with 10 ng/ml fresh IL-2 for 2 hr prior to harvesting cell pellets. For Western blot analysis of JAK inhibitor–treated cell lines, cells were exposed to increasing concentrations of JAK inhibitors (upadacitinib or tofacitinib) or DMSO vehicle for 4 hr. During this period, cells were starved of IL-2 for the first 3.5 hr, followed by 10 ng/ml IL-2 restimulation for the final 30 min. For Western blotting of primary mouse splenocytes, erythrocyte lysis was performed prior to collection of splenocyte pellets. 30 µg total protein was subjected to immunoblotting using standard techniques.

Nitrocellulose membranes (0.45 µm Cytiva Amersham Protran, Fisher Scientific) were blocked in Odyssey Blocking Buffer (Intercept TBS Blocking Buffer, LI-COR Biosciences) and incubated with the respective antibody diluted in the same buffer. Details of the antibodies used for Western blotting are available in **Supplementary Table 2**. Images were obtained using IRDye fluorescent secondary antibodies and an Odyssey CLx imaging system (LI-COR Biosciences).

**Cell competition assay**

For CRISPR/Cas9-mediated knockout of the human $STAT5B^{N642H}$ transgene, C15 cells were lentivirally transduced with a Cas9- and GFP-expressing LentiCRISPRv2GFP vector (#82416; Addgene) additionally containing human *STAT5B* single guide RNAs (sgRNAs) or non-targeting control sgRNAs (sequences listed in **Supplementary Table 3**). The population of successfully transduced cells (percentage of $GFP^+$ cells) were monitored over time using an iQue3 flow cytometer (BioScience, Sartorius Group) and normalised to day 3 post transduction and to the non-targeting controls.

**Sanger Sequencing**

Cells were harvested and genomic DNA was isolated using standard protocols. DNA concentration and purity were determined using a NanoDrop 2000 spectrophotometer (Thermo Fisher Scientific). A sequence encoding the STAT5B C-terminal region was amplified by PCR. The 646 bp product was separated by agarose gel electrophoresis and isolated using a MinElute PCR Purification Kit (Qiagen). Sanger sequencing was carried out by Microsynth Austria GmbH. All primer sequences are listed in **Supplementary Table 4**.

**T-cell receptor rearrangement**

For assessing γ and δ TCR rearrangement, standard PCR mix composed of 40 ng isolated genomic DNA, 250 nM forward and reverse specific primers (**Supplementary Table 5**), 1x buffer, 200 µM dNTPs, 1.5 mM $MgCl_2$, 0.1 µl of Taq DNA polymerase, and nuclease-free water to a total of 20 µl was subjected to 35 cycles of PCR with the following conditions. For *Vγ4, Vγ5, Vγ6, Vγ7* chains: 15 seconds at 95°C (denaturation), 15 seconds at 56°C (annealing), 30 seconds at 72°C (extension) and 10 minutes at 72°C. For *Vγ1, Vγ2, Vδ4, Vδ5* and *Gapdh,* an annealing temperature of 55°C was instead used. PCR products were subjected to standard agarose gel electrophoresis.

**Proliferation assay**

C2, C6 and C15 cells were seeded at $1x10^4$ cells/well, and DERL-2 and DERL-7 cells were seeded at $2x10^4$ cells/well, into 24-well plates. Each cell line was seeded in the presence or absence of 10 ng/ml IL-2 and in technical triplicates for each time point. At the respective time points, cells were harvested into FACS tubes, washed once with PBS, and cell numbers were quantified with the addition of Precision Counting Beads (Biolegend) using a BD FACSCanto II flow cytometer, with FACSDiva (BD Biosciences) and FlowJo (version 10.5.3) software.

**Cytology**

For murine γδ T-cell lines, coverslips were placed at the bottom of each well in a 24-well tissue culture plate. $2x10^6$ cells in 500 µl 1x PBS were seeded per well onto the coverslips. After 30 min at room temperature, the PBS was carefully aspirated. As a control, primary γδ T cells were isolated from lymph nodes and spleens of WT C57BL/6N mice by FACS sorting (Thy1.2$^+$, TCRδ$^+$, TCRβ$^-$; see **Supplementary Table 6**), and $1.5x10^5$ cells were resuspended in 200 µl PBS. The primary γδ T cells were centrifuged onto a glass slide for 3 min at 800 rpm using a Cytospin3 centrifuge (Shandon), and then left to air dry at room temperature. All cells were stained with HAEMA Quick-stain (DIFF Quick, Labor und Technik Eberhard Lehmann GmbH, Germany), according to the manufacturer's protocol. A BX 53 LED light microscope fitted with an Olympus SC50 camera was used to image the cell morphology.

**RNA sequencing**

For murine samples: C2, C6 and C15 cell lines were cultured in biological triplicates and harvested by trypsinization. Primary murine γδ T cells from three individual female WT C57BL/6N mice were isolated using lymph node single cell suspensions. Murine cells were stained with FACS antibodies to discriminate viability dye$^-$ Ter119$^-$ TCRβ$^-$ CD45.2$^+$ TCRδ$^+$ cells (see **Supplementary Table 6**). 500 cells/well were collected from all murine samples by FACS sorting, using a FACSAria III cell sorter, into a hard-shell, low-profile, thin-wall 96-well skirted PCR plate (Bio-Rad) containing 4 µl lysis buffer per well (2 U/µl RNase inhibitor (Clontech) in 0.2% (v/v) Triton X-100). Plates were kept at -80°C until further processing at the Biomedical Sequencing Facility (BSF; CeMM, Vienna, Austria). For preparing NGS libraries, we followed the Smart-seq2 protocol.[2] The subsequent library preparation from the amplified cDNA was performed using a Nextera XT DNA library prep kit (Illumina, San Diego, CA, USA). Library concentrations were quantified with a Qubit 2.0 Fluorometric Quantitation system (Life Technologies, Carlsbad, CA, USA) and the size distribution was assessed using a 2100 Bioanalyzer instrument (Agilent, Santa Clara, CA, USA). For sequencing, samples were diluted and pooled into NGS libraries in equimolar amounts and sequenced on a HiSeq 4000 instrument (Illumina) in 50 bp single end mode.

For human samples: EDTA peripheral blood mononuclear cells (PBMCs) of healthy human donors were isolated by density gradient centrifugation (#25-072-CV, Corning). $CD3^+$ T-cell enrichment from PBMCs was obtained using magnetic cell separation according to the manufacturer's instructions (negative selection, #480021, Biolegend). DERL-2 and DERL-7 cell lines were cultured in biological triplicates, harvested, and RNA extraction was performed with an RNeasy Plus Micro kit (Qiagen) according to the manufacturer's protocol. RNA concentration and purity were measured with a NanoDrop 2000 spectrophotometer. RNA was subjected to polyA-based library preparation and sequenced on a NovaSeq 6000 platform (Illumina) according to the manufacturer's instructions.

PRINSEQ-lite[3] (version 0.20.4) was used for data quality filtering, trimming, and length filtering. High-quality reads were aligned using STAR[4] (version 2.7.9a) to the mouse (mm10) reference genome for the Smart-seq2 datasets and to the human (GRCh38) reference genome for RNA-seq datasets. Aligned reads were subsequently processed with Samtools[5,6] (version 1.13). FeatureCounts[7] from the subread package (version 2.0.3) was used for counting reads per gene. Normalization and differential expression analysis were conducted using DESeq2.[8] For visualizing the results, ggplot2[9] was used in R to create heatmaps. To facilitate cross-species analysis, Bioconductor libraries were used, including Orthology.eg.db,[10] org.Mm.eg.db[11] and org.Hs.eg.db,[12] for translating mouse gene identifiers to their human orthologs.

For patient HSTCL samples, publicly available RNA-seq data from the Gene Expression Omnibus dataset GSE57944[13] were extracted, including case 4, case 5, case 7 and normal spleen samples. These data were processed following the same pipeline described above for human RNA-seq data. To generate the combined heatmap, the ComplexHeatmap[14] package was applied using Z-scores of normalized expression values from all three datasets. The depicted gene list was obtained by identifying significantly differentially expressed genes (P-adj ≤ 0.05 and log2 fold change > 2 or < -2) between the three patient samples and the healthy human controls that had one-to-one orthologs in mouse and were also differentially regulated in the other two comparisons. Gene set enrichment analysis (GSEA)[15] was performed using these overlapping DEGs, pre-ranked by log2 fold change values, and gene sets obtained from the Human Collection Molecular Signatures Database (MSigDB), using GSEA software v4.4.0 (Broad Institute) with 1000 permutations. The enriched pathways were then visualized as a bubble plot with the size of each bubble representing -log10 False Discovery Rate (FDR), and the colour corresponding to either up- or down-regulated pathways, using the Google Colaboratory platform. Additionally, GSEA was performed with normalized gene expression count lists from the murine and human cell lines and respective controls, obtained from the DESeq2 analysis, and gene sets (equivalent mouse and human collections) obtained from the MSigDB, using GSEA software v4.3.2 (Broad Institute) with 1000 permutations and ‘gene_set’ as the permutation type.

**Cell killing assay**

Murine YAC-1 lymphoma cells, used as target cells, were labelled with CSFE (Invitrogen) and $1x10^5$ cells were seeded per well into flat bottom 96-well plates. The clonal murine γδ T-cell lines were then added at the indicated effector:target cell ratios. Primary mouse NK cells, used as a positive control, were isolated and cultured in media supplemented with 3000 U/ml human IL-2 (Novartis), as previously described.[16] The plates were centrifuged at 11 *g* for 2 min to bring the cells into close proximity. After 4 hr incubation at 37°C, the specific lysis was assessed by flow cytometry using SYTOX$^{TM}$ Blue Dead Cell Stain (Invitrogen) to quantify lysed target cells. Percentage of specific lysis was calculated as follows: [% SYTOX+ CFSE+ cells after co-incubation with effector cells] – [% SYTOX+ CFSE+ cells without addition of effector cells (spontaneous lysis control)].

**Mouse strains and housing**

C57BL/6 Ly5.1 (B6.SJL-Ptprc$^{a}$Pepc$^{b}$/BoyCrl), *NOD-scid IL2Rgamma$^{null}$* (NSG) and *NOD-scid IL2Rgamma$^{null}$-3/GM/SF* (NSGS) mice were obtained via in-house breeding at the University of Veterinary Medicine Vienna (Vienna, Austria). Additionally, NSG mice were purchased from Janvier Labs. Female mice between 8-12 weeks of age were used for the experiments in this study. Mice were maintained under specific pathogen-free (SPF) conditions in individually ventilated cages at the University of Veterinary Medicine Vienna. Mice were kept in a 12/12-hour light/dark cycle and received standard food and water *ad libitum*.

***In vivo* allograft models and JAK inhibitor treatment**

Post C15 or M2 tumor cell transplantation via tail vein injection, recipient mice were monitored daily for physical signs of disease (hunching, slow movement, ruffled fur). The percentage of peripheral blood tumor cells, obtained from puncture of the *vena facialis*, was determined every two weeks via FACS analysis of CD45.2$^{+}$ cells (in Ly5.1/CD45.1 recipients) or FLAG-tag$^{+}$ cells (NSG recipients). Mice were considered to have reached end-stage disease when physical signs of disease reached ethical limits and/or the percentage of tumor cells in the blood reached 25%, at which point mice were humanely sacrificed. Directly prior, whole blood samples were collected via heart puncture into EDTA-treated tubes (MiniCollect® K3EDTA, Greiner Bio-One). Immediately following euthanasia, mice were subjected to whole-body perfusion with PBS to remove blood contamination from tissues.

For *in vivo* upadacitinib treatment, C15 cells were intravenously transplanted into 15 female NSG mice, aged 11-12 weeks, as described above. Ten days post transplantation, recipient mice were randomly distributed into two groups and treatment commenced of once-daily oral gavage with either 10 mg/kg upadacitinib (MedChemExpress; $n$ = 8) or vehicle (5% DMSO, 40% PEG-300, 5% Tween-80, 50% PBS; $n$ = 7). Mice were treated five days per week for four weeks (cycle 1) followed by another two weeks of treatment (cycle 2), with a one-week break in treatment between cycles. Once any of the mice in the experiment reached end-stage disease (as determined by our termination criteria in accordance with ethical limits), all mice were sacrificed as described above and comparative end-point analyses were conducted.

**Preparation of cells from mouse tissues**

Single-cell suspensions were prepared from spleen, lymph nodes and liver by crushing organs in ice-cold PBS through a 100 µm cell strainer (BD Biosciences). Bone marrow cells were harvested from the femur and tibia by cutting one end of the bone and centrifuging the contents at 2500 x $g$ for 1 min into an Eppendorf tube containing plain RPMI media. Erythrocytes from spleen, bone marrow and peripheral blood were lysed using Ammonium-Chloride-Potassium (ACK) buffer (150 mM $NH_4CO_3$, 10 mM $KHCO_3$, 1 mM EDTA, pH 7.2). Liver cells were pelleted by centrifugation and then resuspended in 4 ml 40% Percoll® (Sigma-Aldrich), layered onto 4 ml 70% Percoll®, and centrifuged at 800 x $g$ for 20 min at room temperature with deceleration set to 0. Leukocytes from the interphase were collected and washed once in RPMI 1640 media supplemented with 10% hi-FBS.

**Blood biochemistry and hematocytometry**

Whole blood from mice was collected in EDTA-treated tubes (MiniCollect® K3EDTA, Greiner Bio-One). Platelets, white blood cells (WBCs) and haemoglobin levels were measured from whole blood using an animal blood counter (Scil Vet ABC). To analyse alanine aminotransferase (ALT), aspartate aminotransferase (AST) and blood urea nitrogen (BUN) levels, whole blood was centrifuged at 5000 x $g$ for 5 min at room temperature, plasma was collected and analysed undiluted or at a 1:3 dilution in normal saline (0.9%) using an IDEXX Vet Test 8008 Veterinary Chemistry Analyzer.

**Immunohistochemistry and H&E staining**

Mouse organs were incubated for 24 hr in 4% phosphate-buffered formaldehyde solution (Roti-Histofix; Carl Roth), dehydrated, embedded and cut into 2.5 μm thick sections. Immunohistochemical stainings of CD3, Ki67, FLAG and TCRδ, as well as hematoxylin and eosin (H&E) staining, were performed using standard protocols with the following specifications. Heat-mediated antigen retrieval was performed in citrate buffer at pH 6.0 (Dako) in an autoclave for 20 min (CD3, Ki67), or in a microwave for 7 min at 800 W and 15 min at 290 W (FLAG). CD3 staining (CD3ε rabbit mAb, #85061, Cell Signaling Technology; 1:300) and Ki67 staining (Ki67 rabbit mAb, #12202, Cell Signaling Technology; 1:1000) were performed with Mouse-To-Mouse Blocking Reagent (#MTM125, ScyTek Laboratories), secondary antibodies from an UltraTek HRP Anti-Polyvalent Staining Kit (#AFN600, ScyTek Laboratories) and AEC-Chromogen substrate (#ACD030, ScyTek Laboratories). FLAG staining (FLAG rabbit mAb, #14793, Cell Signaling Technology; 1:500) was performed with 5% goat-serum blocking, secondary antibodies from SignalStain Boost IHC Detection Reagent (HRP rabbit, #8114, Cell Signalling Technology) and ImmPACT DAB substrate kit (#SK-4105, Vector Laboratories). TCRδ staining (TCR δ H-41, #sc-100289, Santa Cruz Biotechnology; 1:100) was performed after heat-mediated antigen retrieval in TRIS-EDTA buffer at pH 9.0 for 20 min (#ZUC029-500, Zytomed Systems) and blocking with Mouse-To-Mouse Blocking Reagent (#MTM125, ScyTek Laboratories). Secondary antibody (#POLHRP-100, Zytomed Systems) and visualization (#DAB530, Zytomed Systems) were applied according to the manufacturer’s instructions. Hematoxylin (Mayer′s hemalum solution, #109249, Sigma-Aldrich) and Eosin G (C.I. 45380, #7089, Carl Roth) were applied according to the manufacturer's protocols. Sections were imaged using an Olympus BX53F2 LED light microscope with an Olympus SC50 camera or were scanned with an Evident SLIDEVIEW research slide scanner VS200 and analysed with OlyVIA software (version 3.4.1, Olympus). Percentage and morphology of tumor cell organ infiltration was assessed using CD3 stained organ sections and scored across three mice per genotype by a trained pathologist in a blinded manner.

Immunohistochemical analyses on human HSTCL tissue samples were performed on 2 µm sections of FFPE tissue using an automated BOND-III Immunostainer (Leica Biosystems). Heat induced epitope retrieval was performed with BOND Epitope Retrieval Solution (solution 2, Leica Biosystems #AR9640) for 20 min. Double immunohistochemical staining for pY-STAT5 and TCRδ was performed in a sequential manner, using primary antibodies against phospho-Stat5 (Tyr694, C11C5, rabbit mAb, Cell Signaling Technology #9359; 1:25) and TCRdelta (H-41, mouse mAb, Santa Cruz Biotechnology #sc-100289; 1:300) incubated for 30 min each. pY-STAT5 was stained first and developed using a BOND Polymer Refine Detection Kit (brown color; Leica Biosystems #DS9800), followed by staining for TCRδ and developing with a BOND Polymer Refine Red Detection Kit (magenta color; Leica Biosystems #DS9390), according to the manufacturer's guidelines. Tissue sections were then counterstained with hematoxylin.

**Flow cytometry**

For flow cytometry analysis, cells were transferred to a V-bottom 96-well plate. Fc receptor was blocked with blocking antibody (diluted at 1:100) for 30 min in the dark, and cells were then incubated with antibodies against surface proteins (diluted at 1:200) for 1 hr at 4°C in the dark (**Supplementary Table 6**). For intracellular staining, cells were resuspended in 100 µl of Fix/Perm solution (BD Biosciences) per well in a V-bottom 96-well plate for 20 min at 4°C. Subsequently, cells were washed twice in Perm/Wash Buffer (BD Biosciences), blocked with Fc receptor blocking antibody (diluted at 1:100) and then incubated with antibodies (diluted at 1:200; **Supplementary Table 6**) in Perm/Wash Buffer following the same staining procedure as above. All analyses were performed on a BD FACSCanto II using FACSDiva (BD Biosciences) and FlowJo (version 10.5.3) software or on a Beckman Coulter Life Sciences CytoFLEX benchtop flow cytometer and CytExpert (version 2.4.0.28) software.

***STAT5B* SNP genotyping using digital PCR (dPCR)**

FFPE spleen or bone marrow tissue from five HSTCL patient tumors were cut into 10 µm sections, and 1-5 sections per tumor were collected into a 1.5 ml Eppendorf tube. Genomic

DNA isolation was performed using an AllPrep® DNA/RNA FFPE isolation kit (QIAGEN), according to the manufacturer's protocol. For the deparaffinization step, the protocol using heptane and methanol was used and was performed twice.

dPCR quantification was performed on the 3-colour naica® system (Stilla Technologies, Villejuif, France). The target of interest was a SNP causing the oncogenic somatic missense mutation N642H, located in exon 16 of human *STAT5B* (NM_012448.4: c.A1924C). Assay oligonucleotides were designed using Primer3 (version 2.5.0), integrated into NCBI's Primer-BLAST,[17] and the DNA sequence surrounding the SNP of interest (GenBank: NM_012448.4). To test for cross amplification, the RefSeq Representative Genome Database was restricted to *Homo sapiens*. Six locked nucleic-acid (LNA) monomers were introduced to increase structural stability of the SNP-specific probe. To enhance the disruptive effect of the single mismatch, the probe sequence was shortened to ten nucleotides and designed to melt at a temperature that was 5 to 7°C above the annealing/extension temperature of 60°C.[18] The melting temperature (*Tm*) was predicted using the *Tm* Prediction tool for LNA-enhanced oligonucleotides (Qiagen; https://geneglobe.qiagen.com/us/tools/tm-prediction). LNA monomers were inserted at the polymorphic site located in the probe centre and across the sequence body except for the 5′ terminal base. This allowed cleavage by the 5′ to 3′ nuclease activity of *Taq* DNA polymerase.[19] The SNP genotyping assay used the primers 5'-TGA TTG TTC TGT TTA TTG ATC TAG AGG and 5'-AAT GGA GAA GTC TCT GGT GGT AAA together with the Affinity Plus® hydrolysis probe 5'-FAM-CA+G+A+T+G+C+CAA/TAO™/A-Iowa Black™ RQ, where the substitution is underlined and the "+" sign precedes an LNA base. The number of haploid genomes was counted using an 87-bp-amplicon assay targeting the single-copy gene *RPP30*,[20] with fluorescence generation achieved by a Cy5-labeled double-quenched probe incorporating the internal TAO™ Quencher and the 3' Iowa Black RQ®. Oligonucleotides were synthesised at Integrated DNA Technologies (Leuven, Belgium). The 25 µl reaction volume consisted of 10× naica® Multiplex PCR Mix (2.5 µl buffer A and 1 µl buffer B; Stilla Technologies), 800 nM of each primer, 300 nM of each probe and 2 µl DNA. The reaction mixture was loaded on a Sapphire chip capable of forming up to 30,000 droplets per sample. The chip was placed into the Geode instrument for partitioning the reaction into droplets, formation of droplet crystals and amplification. Amplification was performed according to the following parameters: initial denaturation step at 95°C for 3 min, 45 cycles at 95°C for 10 sec and 60°C for 40 sec. Fluorescence images were obtained using the naica® Prism3 fluorescence reader. Spill-over compensation was performed manually and was applied before data analysis with the Crystal Miner™ software version 4.0.10.3 (Stilla Technologies).

Accuracy of the genotyping results was controlled through the use of genomic DNA from human cell lines with known *STAT5B* wild-type (WT) or N642H mutant alleles; the WT sequence of *STAT5B* was controlled for using Karpas 384 cells, and homo- or heterozygous genotypes were represented by the DERL-7 and DERL-2 cell lines, respectively. To determine the sensitivity of the dPCR assay targeting $STAT5B^{N642H}$, genomic DNA from Karpas 384 (WT *STAT5B*) and DERL-7 (homozygous $STAT5B^{N642H}$) cells was normalized to the same concentration (2,620 copies/µl) by measuring the concentration of single copy gene *RPP30* by dPCR. Karpas 384 cell DNA was then spiked with 0%, 1%, 5% or 10% DNA from DERL-7 cells, and $STAT5B^{N642H}$ and *RPP30* copy numbers were quantified using dPCR. $STAT5B^{N642H}$ variant allele frequency (VAF) was then calculated from the measured $STAT5B^{N642H}$ copy numbers as a percentage of *RPP30* copy numbers from each sample.

**Statistics**

GraphPad Prism v8.4.3 software was used for statistical analyses, applying unpaired two-tailed Student's t-tests for comparison of two groups, and two-way ANOVA with Tukey post-test for comparison of multiple groups with a time variable. *$p < 0.05$, **$p < 0.01$, ***$p < 0.001$, ****$p < 0.0001$.

**Graphical licences**

The cell line generation schematic was created using BioRender.com, under the agreement number KJ26MEJQCR.

**Supplementary Tables**

**Supplementary Table 1: HSTCL Patient Samples**

| Patient # | Age | Sex | Tissue (FFPE) | Status of sample (FFPE) | Tissue (cryopreserved, viable) | Status of sample (cryopreserved, viable) | Tumor cell % (cryopreserved samples) | Healthy cell marker | Tumor cell markers |
|---|---|---|---|---|---|---|---|---|---|
| 1 | 37 | M | Bone marrow | Primary diagnosis | Peripheral blood | Relapse (splenectomy-SMILE-bortezomib) | 15% | CD5$^{+}$ | CD3$^{+}$ CD56$^{+}$ TCRδ$^{+}$ |
| 2 | 75 | F | Spleen | Primary diagnosis | Spleen | Primary diagnosis | 5-10% | CD20$^{+}$ | CD2$^{+}$ CD3$^{+}$ CD7$^{+}$ |
| 3 | 32 | M | Spleen | Primary diagnosis, undergoing SMILE protocol (+21d) | Bone marrow | Relapse (splenectomy-SMILE-cyclophosphamide+mitoxantrone+prednisolone+alemtuzumab) | 45% | CD20$^{+}$ | CD2$^{+}$ CD3$^{+}$ CD7$^{+}$ |
| 4 | 30 | M | Spleen | Primary diagnosis | Spleen | Primary diagnosis | 15% | CD20$^{+}$ | CD2$^{+}$ CD56$^{+}$ |
| 5 | 50 | M | Spleen | Primary diagnosis | Spleen | Primary diagnosis | 5-10% | CD20$^{+}$ | CD2$^{+}$ CD3$^{+}$ CD7$^{+}$ |

**Supplementary Table 2: Western Blot Antibodies**

| Antibody | Company | Cat. # | Dilution |
|---|---|---|---|
| DYKDDDDK FLAG Tag | Cell Signaling Technology | 14793S | 1:1000 |
| pY-STAT5 | Cell Signaling Technology | 9314S/9351S | 1:1000 |
| STAT5 | BD Biosciences | 610191 | 1:1000 |
| Actin (C11) | Santa Cruz Biotechnology | sc-1615R | 1:10000 |
| β-actin (C4) | Santa Cruz Biotechnology | sc-47778 | 1:5000 |
| HSC70 | Santa Cruz Biotechnology | sc-7298 | 1:5000 |
| IRDye 680RD Goat anti-Mouse IgG | LI-COR | 926-68070 | 1:10000 |
| IRDye 800CW Goat anti-Mouse IgG | LI-COR | 926-32210 | 1:10000 |
| IRDye 680RD Goat anti-Rabbit IgG | LI-COR | 925-68071 | 1:10000 |
| IRDye 800CW Goat anti-Rabbit IgG | LI-COR | 926-32211 | 1:10000 |

**Supplementary Table 3: CRISPR/Cas9 Guide RNAs**

| Target | Forward sequence | Reverse sequence |
|---|---|---|
| hSTAT5B_1 | CACCGTAACGCTTGCATCTGATGAA | AAACTTCATCAGATGCAAGCGTTAC |
| hSTAT5B_2 | CACCGCCTCAAACGTCTGGTTGATC | AAACGATCAACCAGACGTTTGAGGC |
| non-targeting control_1 | CACCGCGCTTCCGCGGCCCGTTCAA | AAACTTGAACGGGCCGCGGAAGCGC |
| non-targeting control_2 | CACCGATCGTTTCCGCTTAAGGCG | AAACCGCCGTTAAGCGGAAACGATC |

**Supplementary Table 4: Sanger Sequencing Primers**

<u>PCR Primers</u>

| Target | Forward sequence | Reverse sequence |
|---|---|---|
| *STAT5B* | GGCAATGGTTTGACGGTG | GGATCCACTGACTGTCCATT |

Sequencing Primer

| Target | Sequence |
|---|---|
| *STAT5B* | GCCTCATTGGAATGATGG |

**Supplementary Table 5: TCRγδ Rearrangement PCR Primers**

Primer sequences were designed as previously described.[21] The nomenclature used for mouse γδ T-cell receptor genes is based on the Heilig and Tonegawa's system.[22]

| Target | Forward sequence | Reverse sequence |
|---|---|---|
| *Vγ1* | CTTCCATATTTCTCCAACACAGC | ACTACGAGCTTTGTCCCTTTGG |
| *Vγ2* | | ACTATGAGCTTTGTTCCTTCTGCAA |
| *Vγ4* | TGGACATGGGAAGTTGGAG | CAGAGGGAATTACTATGAGC |
| *Vγ5* | GATCAGCTCTCCTTTACCC | |
| *Vγ6* | CTGGGGTCATATGTCATCAA | |
| *Vγ7* | GCTAACCTACCATTCTCTGT | |
| *Vδ4* | CCGCTTCTCTGTGAACTTCC | CAGTCACTTGGGTTCCTTGTCC |
| *Vδ5* | CAGATCCTTCCAGTTCATCC | |
| *Gapdh* | AGGTCGGTGTGAACGGATTTG | TGTAGACCATGTAGTTGAGGTCA |

**Supplementary Table 6: Flow Cytometry Antibodies**

| Surface staining - mouse | | | | |
|---|---|---|---|---|
| **Target** | **Fluorochrome** | **Clone** | **Company** | **Cat. #** |
| CD90.2 (Thy1.2) | APC | 53-2.1 | Thermo Fisher Scientific | 17-0902-81 |
| CD90.2 (Thy1.2) | PE/Cy7 | 53-2.1 | Biolegend | 140309 |
| CD90.2 (Thy1.2) | eFluor450 | 53-2.1 | Thermo Fisher Scientific | 48-0902-82 |
| CD3ε | FITC | 145-2C11 | Biolegend | 100306 |
| CD4 | PE | GK1.5 | Thermo Fisher Scientific | 12-0041-83 |
| CD8α | PerCP-Cy5.5 | 53-6.7 | Thermo Fisher Scientific | 45-0081-82 |

| | | | | |
|---|---|---|---|---|
| CD11b | PerCP-Cy5.5 | M1/70 | Thermo Fisher Scientific | 45-0112-80 |
| NK1.1 | PE/Cy7 | PK136 | Biolegend | 108713 |
| CD25 | PE/Cy7 | PC61.5 | Thermo Fisher Scientific | 17-0251-82 |
| CD45.1 (Ly5.1) | BV650 | A20 | Biolegend | 110735 |
| CD45.1 (Ly5.1) | PE | A20 | Thermo Fisher Scientific | 12-0453-82 |
| CD45.2 (Ly5.2) | PE/Cy5 | 104 | BD Biosciences | 552950 |
| CD45.2 (Ly5.2) | BV605 | 104 | Biolegend | 109841 |
| CD45.2 (Ly5.2) | APC | 104 | Thermo Fisher Scientific | 17-0454-82 |
| CD69 | PE | H1.2F3 | Thermo Fisher Scientific | 11-0691-82 |
| CD117 | FITC | 2B8 | Thermo Fisher Scientific | 15-1171-82 |
| TCRβ | PE/Cy7 | H57-597 | BD Biosciences | 560729 |
| TCRβ | APC | H57-597 | BD Biosciences | 553174 |
| TCRδ | BV421 | GL3 | BD Biosciences | 562892 |
| TCRδ | FITC | GL3 | Thermo Fisher Scientific | 11-5711-85 |
| Ter119 | APC/Cy7 | TER-119 | Biolegend | 116223 |
| FOXP3 | APC | FJK-16s | Thermo Fisher Scientific | 17-5773-80 |
| Rat IgG2a,κ | APC | eBR2a | Thermo Fisher Scientific | 17-4321-81 |
| Armenian hamster IgG | APC | HTK888 | Biolegend | 400911 |
| Rat IgG2a,κ | PE | eBR2a | Thermo Fisher Scientific | 12-4321-81A |
| Rat IgG2b,κ | PE | RTK4530 | Biolegend | 400607 |
| Armenian hamster IgG | FITC | HTK888 | Biolegend | 400905 |
| Rat IgG2a,κ | PerCP-Cy5.5 | eBR2a | Thermo Fisher Scientific | 45-4321-80 |
| Armenian hamster IgG | PE/Cy7 | HTK888 | Biolegend | 400921 |
| Armenian hamster IgG | BV421 | HTK888 | Biolegend | 400935 |
| **Surface staining - human** | | | | |
| CD45 | APC | HI30 | Thermo Fisher Scientific | 17-0459-42 |
| CD3ε | FITC | UCHT1 | Biolegend | 300405 |

| | | | | |
|---|---|---|---|---|
| TCRδ | BV421 | B1 | Biolegend | 331217 |
| TCRδ | PE | 11F2 | BD Biosciences | 333141 |
| TCRαβ | APC | IP26 | Biolegend | 306718 |
| CD2 | PE | TS1/8 | Biolegend | 309208 |
| CD2 | PE | S5.2 | Biolegend | 347405 |
| CD3 | APC/Cy7 | HIT3a | Biolegend | 300318 |
| CD4 | PE | OKT4 | Biolegend | 317410 |
| CD8 | PE | HIT8a | Biolegend | 300908 |
| CD11b | APC | M1/70 | Biolegend | 101212 |
| CD117 | PE/Cy7 | 104D2 | BD Biosciences | 339217 |
| CD25 | APC | BC96 | Biolegend | 302610 |
| CD69 | FITC | FN50 | Biolegend | 310904 |
| CD56 | FITC | HCD56 | Biolegend | 318304 |
| CD56 | APC | NCAM16.2 | BD Biosciences | 341027 |
| CD20 | PE/Cy7 | 2H7 | Biolegend | 302312 |
| CD7 | FITC | CD7-6B7 | Biolegend | 343104 |
| TCRδ | PE | B1 | Biolegend | 331210 |
| CD2 | FITC | RPA-2.10 | Biolegend | 300206 |
| CD5 | PE/Cy7 | L17F12 | Biolegend | 364007 |
| Rat IgG1, κ | APC | RTK2071 | Biolegend | 400411 |
| Mouse IgG2b, κ | FITC | MPC-11 | Biolegend | 400310 |
| Rat IgG2b, κ | PE/Cy7 | RTK4530 | Biolegend | 400617 |
| Mouse IgG2b, κ | PE | MG2b-57 | Biolegend | 401208 |
| **Intracellular staining - mouse** | | | | |
| **Target** | **Fluorochrome** | **Clone** | **Company** | **Cat. #** |
| DYKDDDDK Tag (FLAG) | APC | L5 | Biolegend | 637307 |
| CD3ε | eFluor450 | 145-2C11 | Thermo Fisher Scientific | 48-0031-82 |
| TCRδ | FITC | GL3 | Thermo Fisher Scientific | 11-5711-85 |
| TCRβ | APC | H57-597 | BD Biosciences | 553174 |

| | | | | |
|---|---|---|---|---|
| Armenian hamster IgG | FITC | HTK888 | Biolegend | 400905 |
| **Intracellular staining - human** | | | | |
| CD3ε | FITC | UCHT1 | Biolegend | 300405 |
| TCRδ | PE | 11F2 | BD Biosciences | 347907 |
| TCRαβ | APC | IP26 | Biolegend | 306718 |
| Rat IgG1, κ | APC | RTK2071 | Biolegend | 400411 |
| Mouse IgG2b, κ | PE | MG2b-57 | Biolegend | 401208 |
| **Miscellaneous** | | | | |
| **Target** | **Fluorochrome** | **Clone** | **Company** | **Cat. #** |
| Viability dye | eFluor780 | - | Thermo Fisher Scientific | 65-0865-14 |
| Fc block | - | 93 | Biolegend | 101302 |
| Fc block | - | - | Biolegend | 422301 |
| DAPI | - | - | Biolegend | 422801 |

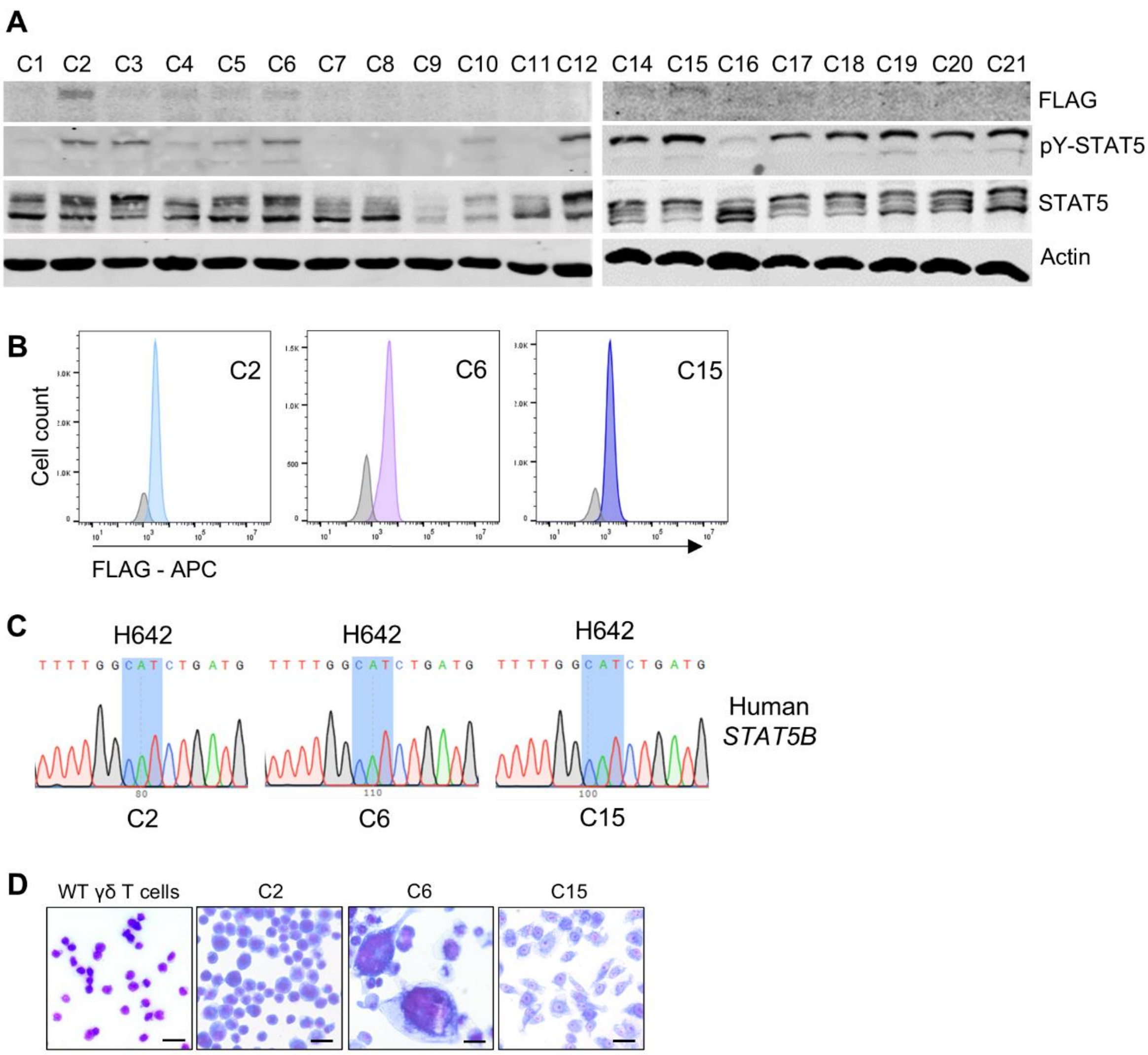


**Supplemental Figure 1. A)** Western blot analyses of FLAG (hSTAT5B$^{N642H}$ transgene), pY-STAT5 and total STAT5 protein levels across 20 clonal murine γδ TCL cell lines, with actin serving as a loading control (*n* = 2). **B)** Representative histograms depicting mean fluorescence intensity of intracellular FLAG (hSTAT5B$^{N642H}$ transgene) protein levels in C2, C6 and C15 cells (blue peaks) compared with isotype control antibodies (grey peaks), as determined by flow cytometry (*n* = 3). **C)** Sanger sequencing of the SH2 domain of human *STAT5B* using genomic DNA isolated from C2, C6 and C15 murine cell lines. The codon 642 is highlighted in blue. **D)** Morphology of C2, C6 and C15 murine γδ TCL cell lines compared with primary murine γδ T cells, analysed by HAEMA Quick-stain and light microscopy at 40x magnification (*n* = 2). Scale bar = 20 µm.

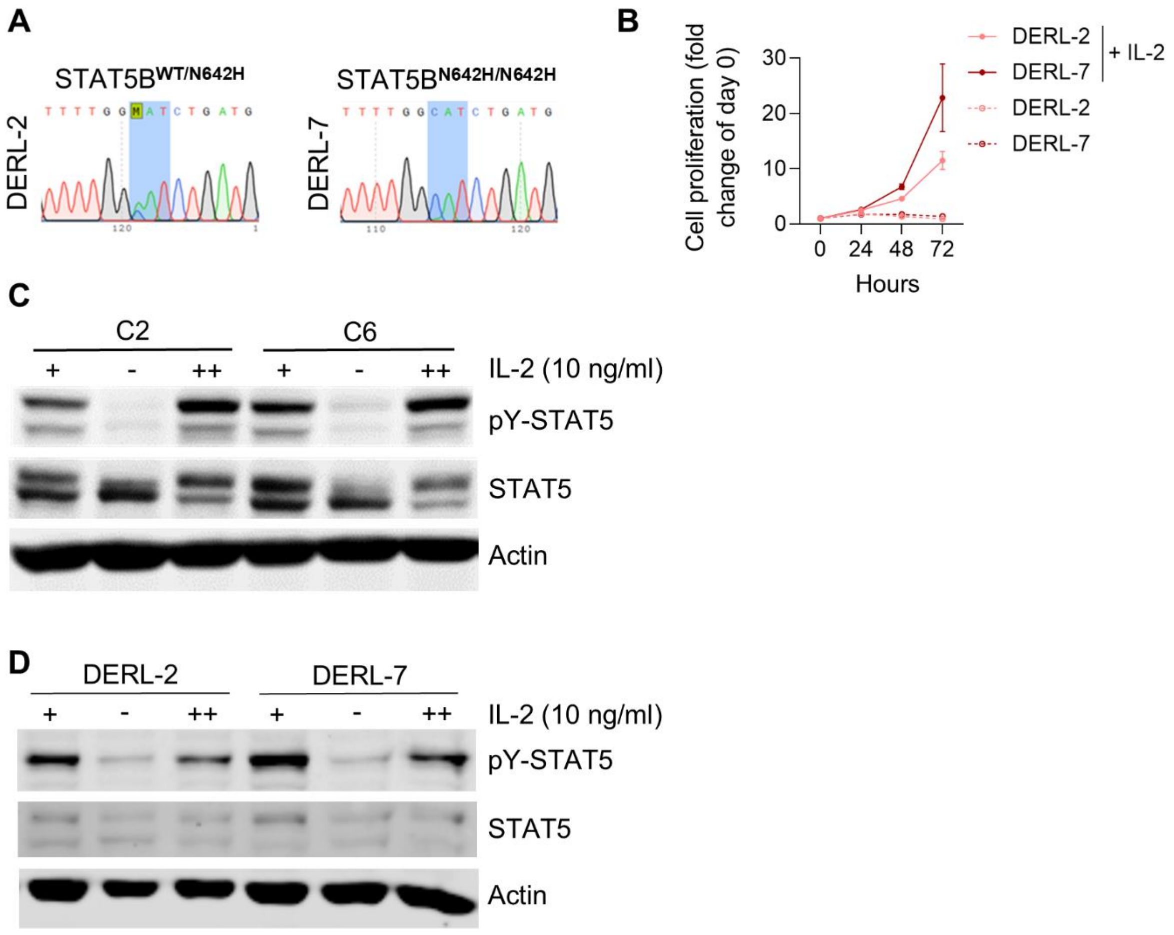


**Supplemental Figure 2. A)** Sanger sequencing of the human *STAT5B* SH2 domain using genomic DNA isolated from DERL-2 and DERL-7 human HSTCL cell lines. The codon for amino acid position 642 is highlighted in blue. **B)** Cell proliferation of human DERL-2 and DERL-7 HSTCL cell lines in the presence or absence of IL-2 over 72 hr, measured by flow cytometry. Data are graphed as mean (± SD) of technical triplicates from one experiment, representative of two independent experiments ($n$ = 2). **C-D)** Western blots showing STAT5 activity in C2 and C6 murine cell lines (C) or DERL-2 and DERL-7 human cell lines (D) cultured in media supplemented with IL-2 for 24 hr (+), starved of IL-2 for 8 hr (-), and then restimulated with IL-2 for 1 hr (++). Immunoblotting for pY-STAT5 and total STAT5 was performed, with actin serving as a loading control (murine cell lines, $n$ = 3; human cell lines, $n$ = 2).

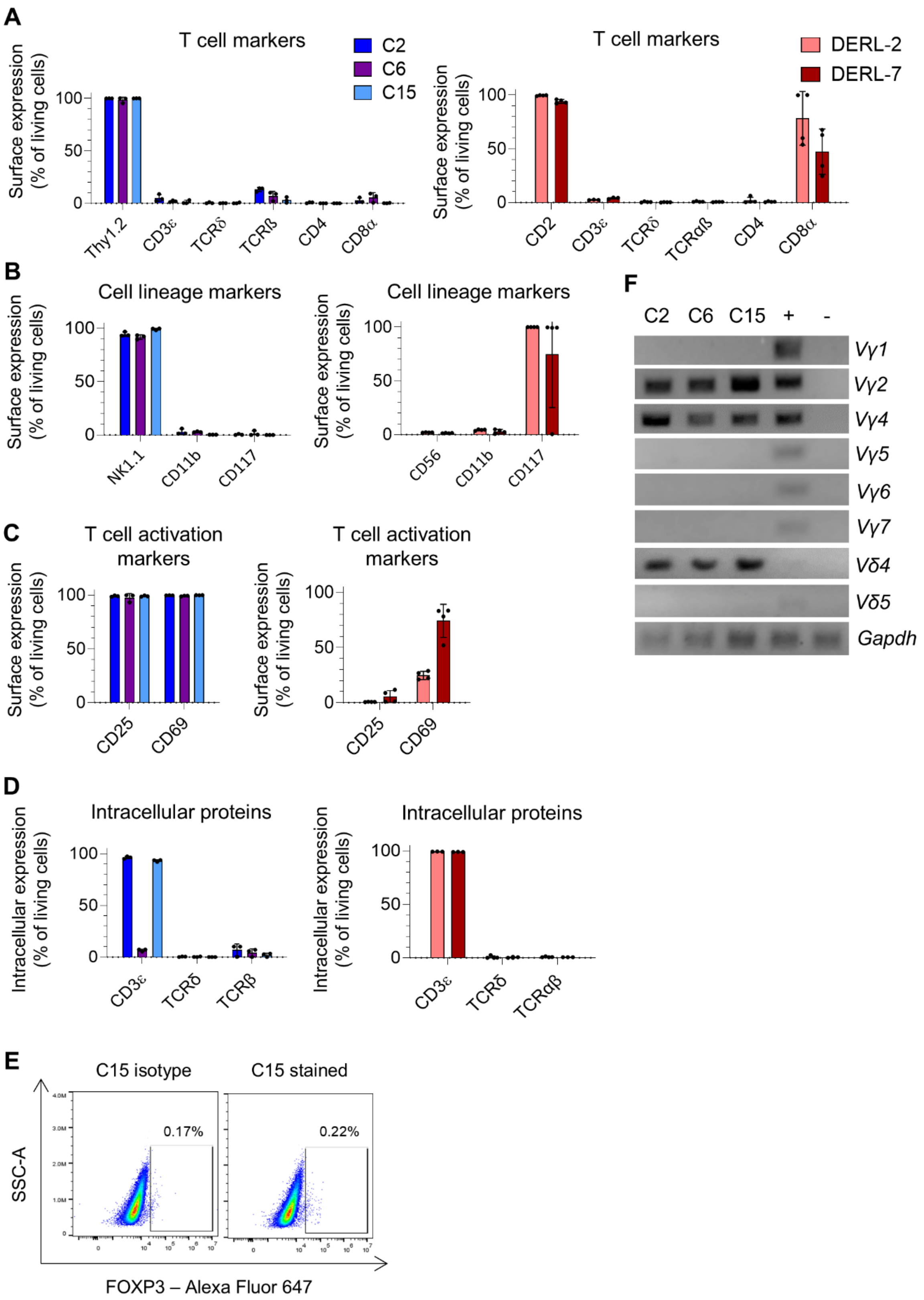


**Supplemental Figure 3. A-D)** Percentage expression of various cell surface T-cell markers (A), cell lineage markers (B) and T-cell activation markers (C), and intracellular proteins (D) on murine C2, C6 and C15 as well as human DERL-2 and DERL-7 cell lines, measured by flow cytometry. Data are graphed as mean (± SD) of biological replicates from three independent experiments (*n* = 3). **E)** Representative FACS plots depicting percentage of C15 cells expressing intracellular FOXP3 protein compared with isotype control staining, determined by flow cytometry (*n* = 3). **F)** T-cell receptor rearrangement analysis of the C2, C6 and C15 cells conducted by PCR, examining murine *Vγ1, Vγ2, Vγ4, Vγ5, Vγ6, Vγ7, Vδ4* and *Vδ5* chains (nomenclature based on the Heilig and Tonegawa's system[22]). *Gapdh* served as an internal control, lymph node cells from *Vav1*-hSTAT5B$^{N642H}$ transgenic mice[1] were used as a positive control, and Ba/F3 cells were used as a negative control (*n* = 2).

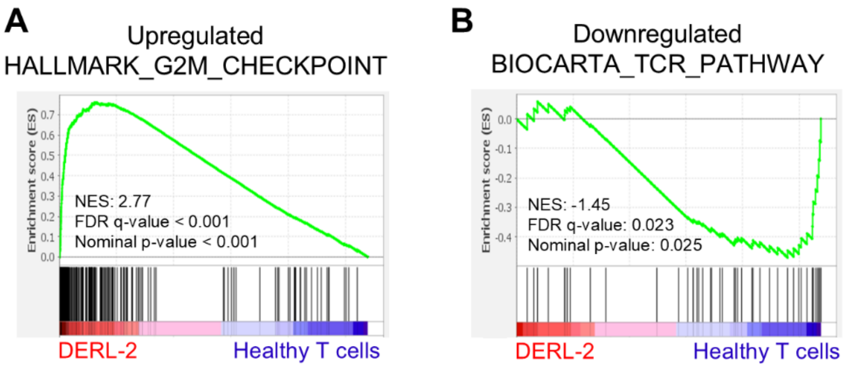


**Supplemental Figure 4. A-B)** GSEA of DEGs from the human DERL-2 cell line compared to healthy control human T cells, displaying significant upregulation of G2/M checkpoint pathway genes (A) and downregulation of TCR pathway genes (B). NES, normalized enrichment score.

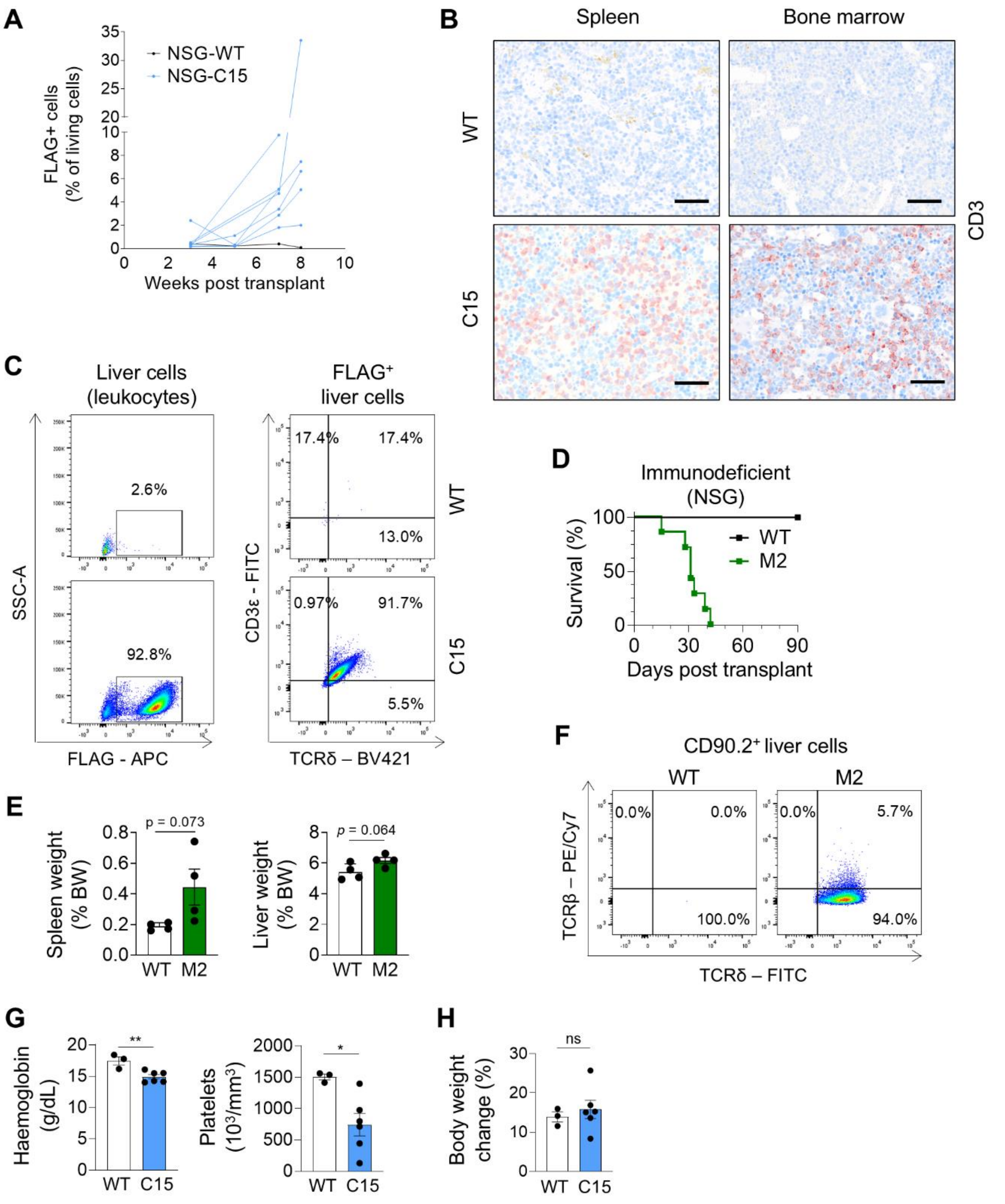


**Supplemental Figure 5. A)** Percentage of FLAG$^{+}$ C15 tumor cells in the peripheral blood of WT ($n$ = 1) or C15-recipient ($n$ = 7) NSG mice at various weeks post-transplant, analyzed by flow cytometry. Each line represents an individual mouse. **B)** Representative images from immunohistochemistry (IHC) analysis of CD3 staining in spleen and bone marrow from WT or C15-recipient NSG mice, imaged by light microscopy (scale bar = 50 µm). **C)** Representative FACS plots showing percentage of FLAG$^{+}$ C15 tumor cells (left), and their expression (%) of cell surface CD3ε and TCRδ (right), in percoll-isolated leukocytes from the liver of WT and

C15-recipient NSG mice. **D)** Kaplan-Meier survival analysis of NSG mice transplanted with M2 cells (*n* = 7) via tail vein injection, or control (WT) mice (*n* = 4). **E)** Spleen and liver weights (as % of body weight, BW) of WT and diseased M2-recipient NSG mice. Data are graphed as mean (± SEM). *p* values determined using an unpaired two-tailed Student's t-test. **F)** Representative FACS plots showing percentage of $CD90.2^+$ cells expressing TCRβ or TCRδ in the liver of WT and M2-recipient NSG mice. **G)** Haemoglobin and platelet levels in whole blood of WT (*n* = 3) and C15-recipient NSG mice (*n* = 6), analysed with an animal blood counter (Scil Vet ABC). Data are graphed as mean (± SEM). $*p < 0.05$, $**p < 0.01$; unpaired two-tailed Student's t-test. **H)** Change in body weight (percentage) of WT (*n* = 3) and C15-recipient NSG mice (*n* = 6) from the date of tumor cell transplantation to the date of end-point analysis. Data are graphed as mean (± SEM). ns = not significant; unpaired two-tailed Student's t-test.

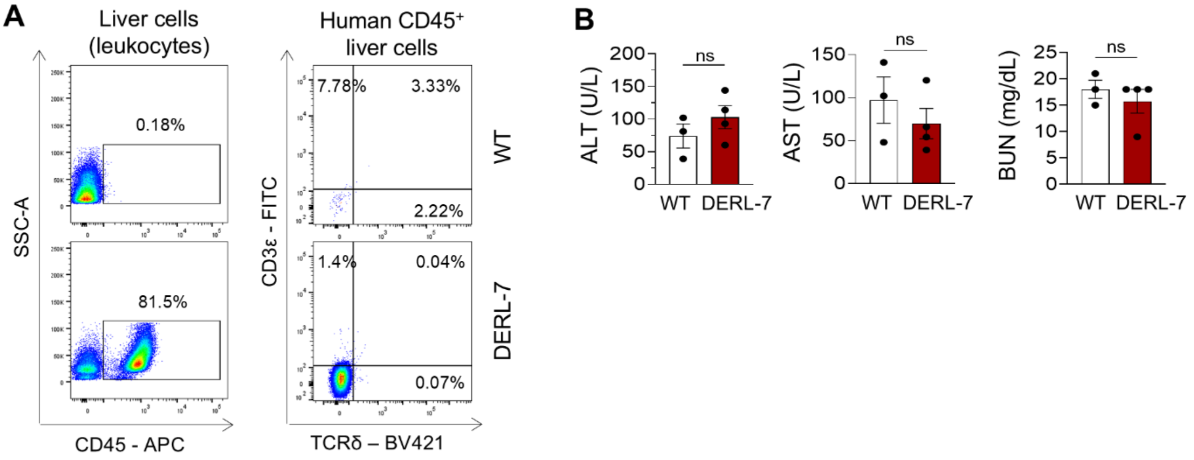


**Supplemental Figure 6. A)** Representative FACS plots showing percentage of human CD45$^+$ DERL-7 tumor cells (left), and their expression (%) of cell surface CD3ε and TCRδ (right), in percoll-isolated leukocytes from the liver of WT (*n* = 3) and DERL-7-recipient (*n* = 5) NSGS mice. **B)** Levels of aspartate aminotransferase (AST), alanine aminotransferase (ALT) and blood urea nitrogen (BUN) in the plasma of WT (*n* = 3) and DERL-7-recipient (*n* = 4) NSGS mice. Data are graphed as mean (± SEM). ns = not significant; unpaired two-tailed Student's t-test.

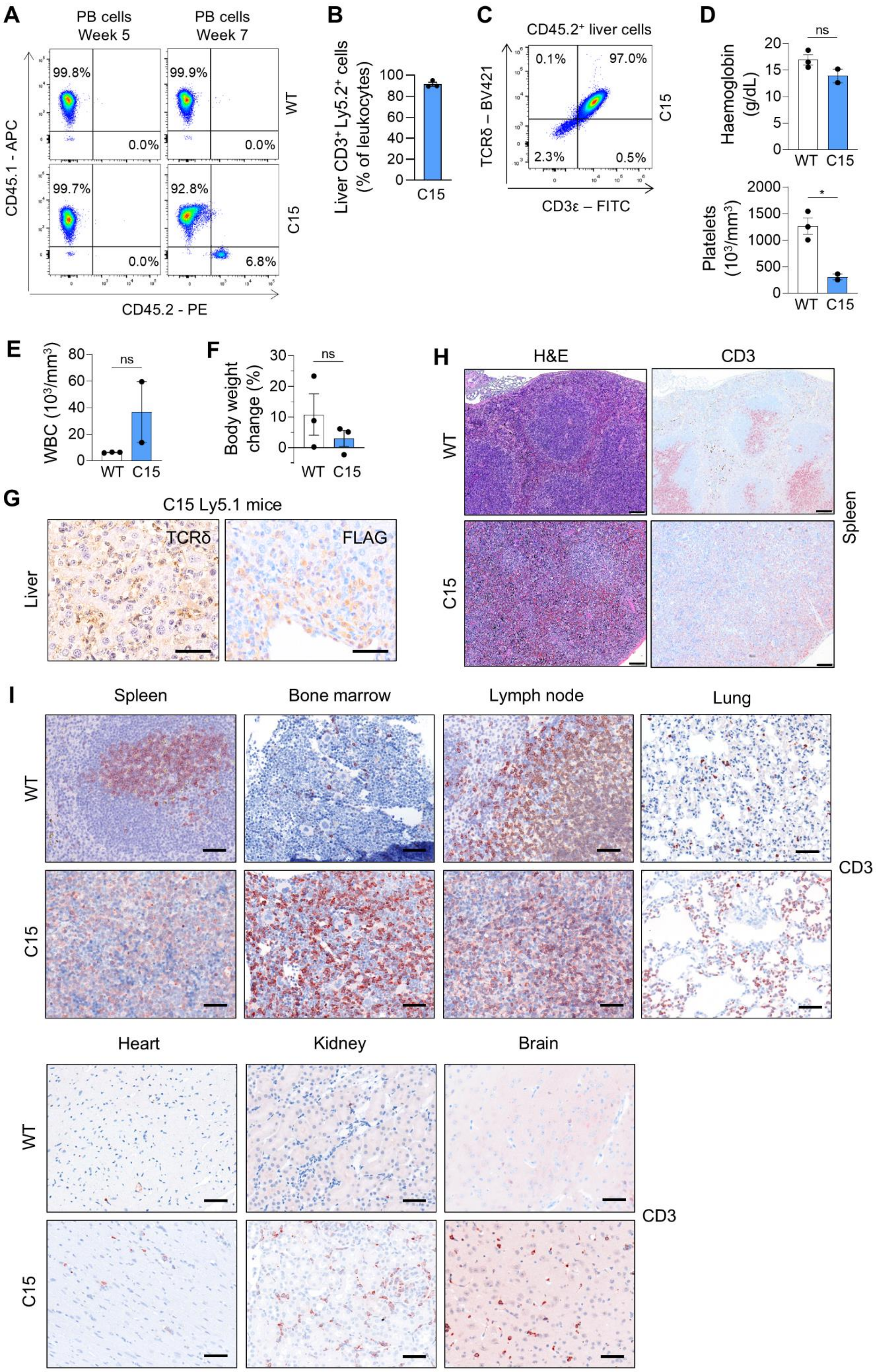
A
PB cells Week 5
PB cells Week 7
99.8%
0.0%
99.9%
0.0%
WT
99.7%
0.0%
92.8%
6.8%
C15
CD45.1 - APC
CD45.2 - PE
B
Liver CD3⁺ Ly5.2⁺ cells (% of leukocytes)
C15
C
CD45.2⁺ liver cells
0.1%
97.0%
2.3%
0.5%
TCRδ – BV421
CD3ε – FITC
C15
D
ns
Haemoglobin (g/dL)
WT C15
*
Platelets (10³/mm³)
WT C15
E
ns
WBC (10³/mm³)
WT C15
F
ns
Body weight change (%)
WT C15
G
C15 Ly5.1 mice
TCRδ
FLAG
Liver
H
H&E
CD3
WT
C15
Spleen
I
Spleen
Bone marrow
Lymph node
Lung
WT
C15
CD3
Heart
Kidney
Brain
WT
C15
CD3

491

**Supplemental Figure 7. A)** Representative FACS plots showing percentages of $CD45.1^+$ and $CD45.2^+$ cells in the peripheral blood (PB) of WT or C15-recipient Ly5.1 mice at 5- and 7-weeks post transplantation. **B)** Quantification of $CD3^+CD45.2^+$ tumor cells (%) in percoll-isolated leukocytes from the liver of C15-recipient Ly5.1 mice ($n$ = 3), using flow cytometry. Data are graphed as mean (± SEM). **C)** Representative FACS plot showing percentages of CD3ε and TCRδ cell surface expression on $CD45.2^+$ liver cells from C15-recipient Ly5.1 mice. **D)** Haemoglobin and platelet levels in whole blood of WT ($n$ = 3) and C15-recipient Ly5.1 mice ($n$ = 2), analysed with an animal blood counter (Scil Vet ABC). Data are graphed as mean (± SEM). *$p$ < 0.05, ns = not significant; unpaired two-tailed Student's t-test. **E)** White blood cell (WBC) counts in the peripheral blood of WT ($n$ = 3) and C15-recipient Ly5.1 mice ($n$ = 2), analysed with an animal blood counter (Scil Vet ABC). Data are graphed as mean (± SEM). ns = not significant; unpaired two-tailed Student's t-test. **F)** Change in body weight (percentage) of WT ($n$ = 3) and C15-recipient Ly5.1 mice ($n$ = 3) from the date of tumor cell transplantation to the date of end-point analysis. Data are graphed as mean (± SEM). ns = not significant; unpaired two-tailed Student's t-test. **G)** Representative images from IHC analysis of TCRδ and FLAG staining in liver from C15-recipient Ly5.1 mice, imaged by light microscopy (scale bar = 50 μm). **H)** Representative images of spleen morphology of WT or C15-recipient Ly5.1 mice using IHC stained for CD3 and H&E, and imaged by light microscopy at 10x magnification (scale bar = 100 μm). **I)** Representative images from IHC analysis of CD3 staining in various organs from WT or C15-recipient Ly5.1 mice, imaged by light microscopy (scale bar = 50 μm).

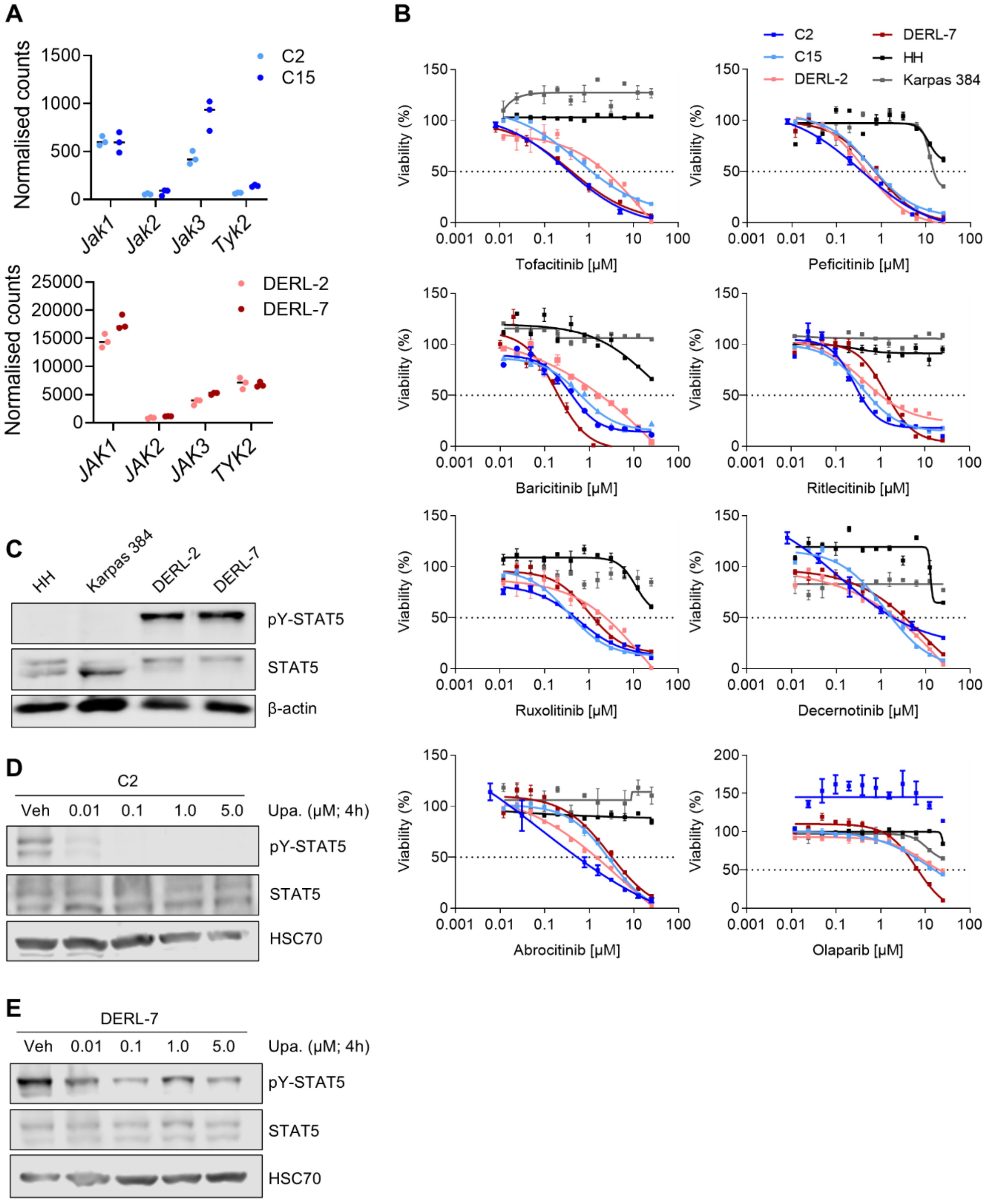


**Supplemental Figure 8. A)** Normalised counts of *Jak1, Jak2, Jak3* and *Tyk2* mRNA in murine C2 and C15 cell lines, and of *JAK1, JAK2, JAK3* and *TYK2* mRNA in human DERL-2 and DERL-7 cell lines, determined by RNA-seq (*n* = 3). **B)** Cell viability curves upon 48 hr treatment of the indicated drugs at various concentrations using murine (C2, C15) and human (DERL-2, DERL-7) HSTCL cell lines, and control human (HH, Karpas 384) TCL cell lines. Data are graphed as mean (± SD) of technical triplicates from one experiment, representative of three independent experiments (*n* = 3). **C)** Western blot showing STAT5 activity in human TCL cell lines (HH, Karpas 384) and HSTCL cell lines (DERL-2, DERL-7). Immunoblotting for pY-STAT5 and total STAT5 was performed, with β-actin serving as a loading control (blots representative of two independent experiments; *n* = 2). **D-E)** Western blots showing STAT5 activity in C2 cells (D) or DERL-7 cells (E) treated with upadacitinib for 4 hr at the indicated concentrations. During this time, cells were starved of IL-2 for 3.5 hr and then restimulated with 10 ng/mL IL-2 for 30 min. Immunoblotting for pY-STAT5 and total STAT5 was performed, with HSC70 serving as a loading control (blots representative of two independent experiments each; *n* = 2). Veh, vehicle (DMSO).

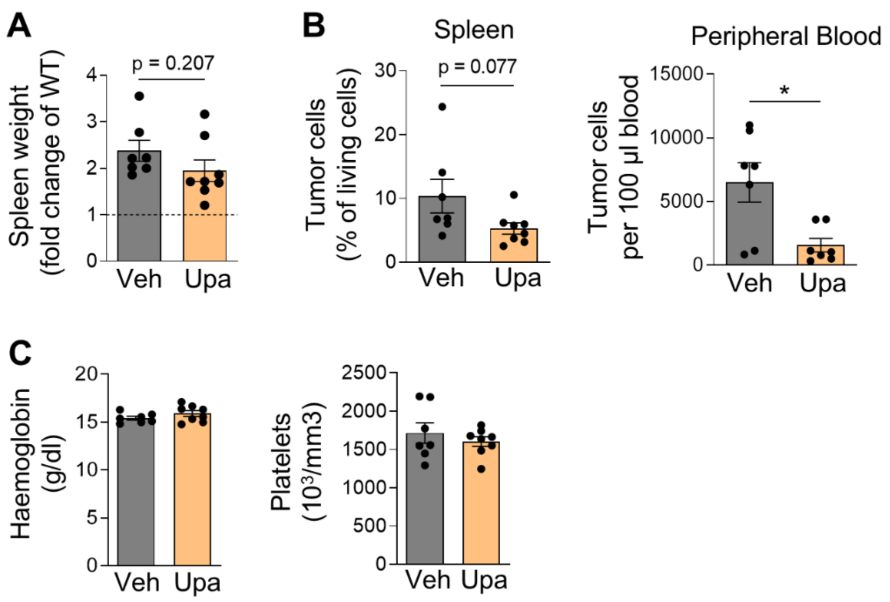


**Supplemental Figure 9. A)** Spleen weights of NSG mice transplanted with C15 cells and treated with vehicle (veh; *n* = 7) or upadacitinib (*n* = 8), graphed as fold change of average spleen weight of WT NSG mice. All mice were analysed 56 days post-transplant. Data are graphed as mean (± SEM); *p* value determined using an unpaired two-tailed Student's t-test. **B)** Quantification of intracellular FLAG+ tumor cell numbers in the spleen and peripheral blood of vehicle or upadacitinib treated C15-recipient NSG mice, analysed by flow cytometry. Data are graphed as mean (± SEM). *$p < 0.05$; unpaired two-tailed Student's t-test. **C)** Haemoglobin and platelet levels in whole blood of vehicle (*n* = 7) or upadacitinib (*n* = 8) treated C15-recipient NSG mice, analysed with an animal blood counter (Scil Vet ABC). Data are graphed as mean (± SEM).

**A**

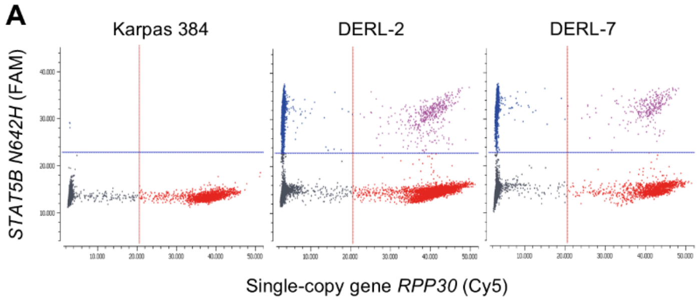


| Cell line | *STAT5B*$^{N642H}$ (cp/µl) | *RPP30* (cp/µl) | *STAT5B*$^{N642H}$ VAF (%) |
|---|---|---|---|
| Karpas 384 | 5 ± 4 | 2189 ± 100 | 0.2 ± 0.2 |
| DERL-2 | 2330 ± 96 | 4259 ± 133 | 54.7 ± 2.3 |
| DERL-7 | 3381 ± 147 | 3119 ± 141 | 108.4 ± 4.7 |

**B**

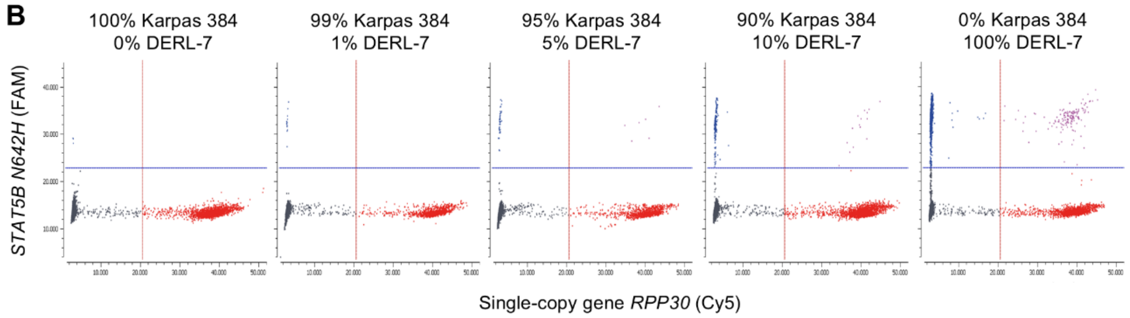


**C**

| % DERL-7 DNA | *STAT5B*$^{N642H}$ (cp/µl) | *RPP30* (cp/µl) | *STAT5B*$^{N642H}$ VAF (%) |
|---|---|---|---|
| 0 | 4.5 ± 4.4 | 2188.8 ± 99.6 | 0.2 ± 0.2 |
| 1 | 20.9 ± 11.4 | 2170.0 ± 118.9 | 1.0 ± 0.5 |
| 5 | 99.5 ± 27.6 | 2186.3 ± 132.7 | 4.6 ± 1.3 |
| 10 | 197.5 ± 27.3 | 2013.8 ± 88.8 | 9.8 ± 1.4 |
| 100 | 2032.5 ± 92.9 | 1788.8 ± 86.8 | 113.6 ± 5.2 |

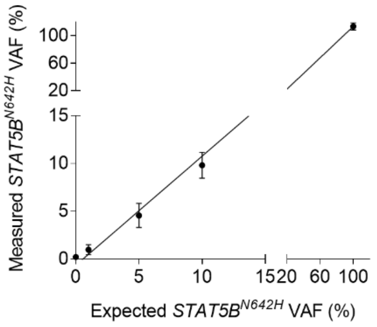


**D**

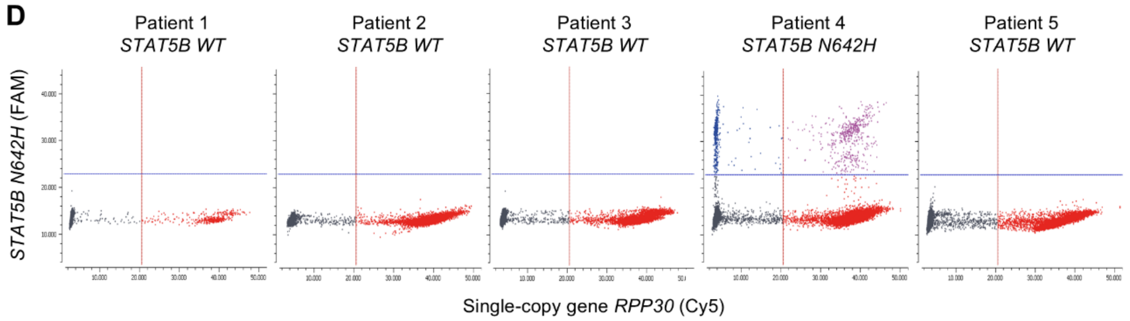


**Supplemental Figure 10. A-C)** Digital PCR (dPCR) variant-specific assay validation for the detection of the *STAT5B*$^{N642H}$ SNP. **A)** Variant allele frequency (VAF) of *STAT5B*$^{N642H}$ measured by dPCR using genomic DNA from Karpas 384 (WT *STAT5B*), DERL-2 (heterozygous *STAT5B*$^{N642H}$), and DERL-7 (homozygous *STAT5B*$^{N642H}$) cells. (*Left*) Data are presented as two-dimensional dPCR scatter plots, with each panel representing a separate measurement of reaction mix containing DNA partitioned into droplets and assessed for *STAT5B*$^{N642H}$ and *RPP30*. The arbitrary threshold distinguishing positive droplets from the background of negative droplets is indicated by a blue (FAM fluorophore) or a red (Cy5 fluorophore) line. (*Right*) Table displaying dPCR data with quantifications of *STAT5B*$^{N642H}$ and *RPP30* copy numbers, and calculation of *STAT5B*$^{N642H}$ variant allele frequency (VAF). Data are presented as mean ± Poisson 95% confidence intervals (as determined by the dPCR software). **B-C)** Measurement of the sensitivity of the dPCR variant-specific assay, using genomic DNA from Karpas 384 cells (WT *STAT5B*) spiked with 0, 1, 5 or 10% DNA from DERL-7 cells (homozygous *STAT5B*$^{N642H}$). *STAT5B*$^{N642H}$ and *RPP30* copy numbers were measured by dPCR. Two-dimensional dPCR scatter plots are shown (B), dPCR data measurements and quantifications of *STAT5B*$^{N642H}$ VAF as percentage of *RPP30* copy number are listed (C, *left*), and the measured *STAT5B*$^{N642H}$ VAFs are graphed against the expected VAFs (C, *right*). Data are presented as mean ± Poisson 95% confidence intervals (as determined by the dPCR software). **D)** Screening for the *STAT5B*$^{N642H}$ variant by dPCR using genomic DNA isolated from FFPE tumor tissues from five HSTCL patients. Two-dimensional dPCR scatter plots are shown, measuring *STAT5B*$^{N642H}$ and *RPP30* copy numbers.

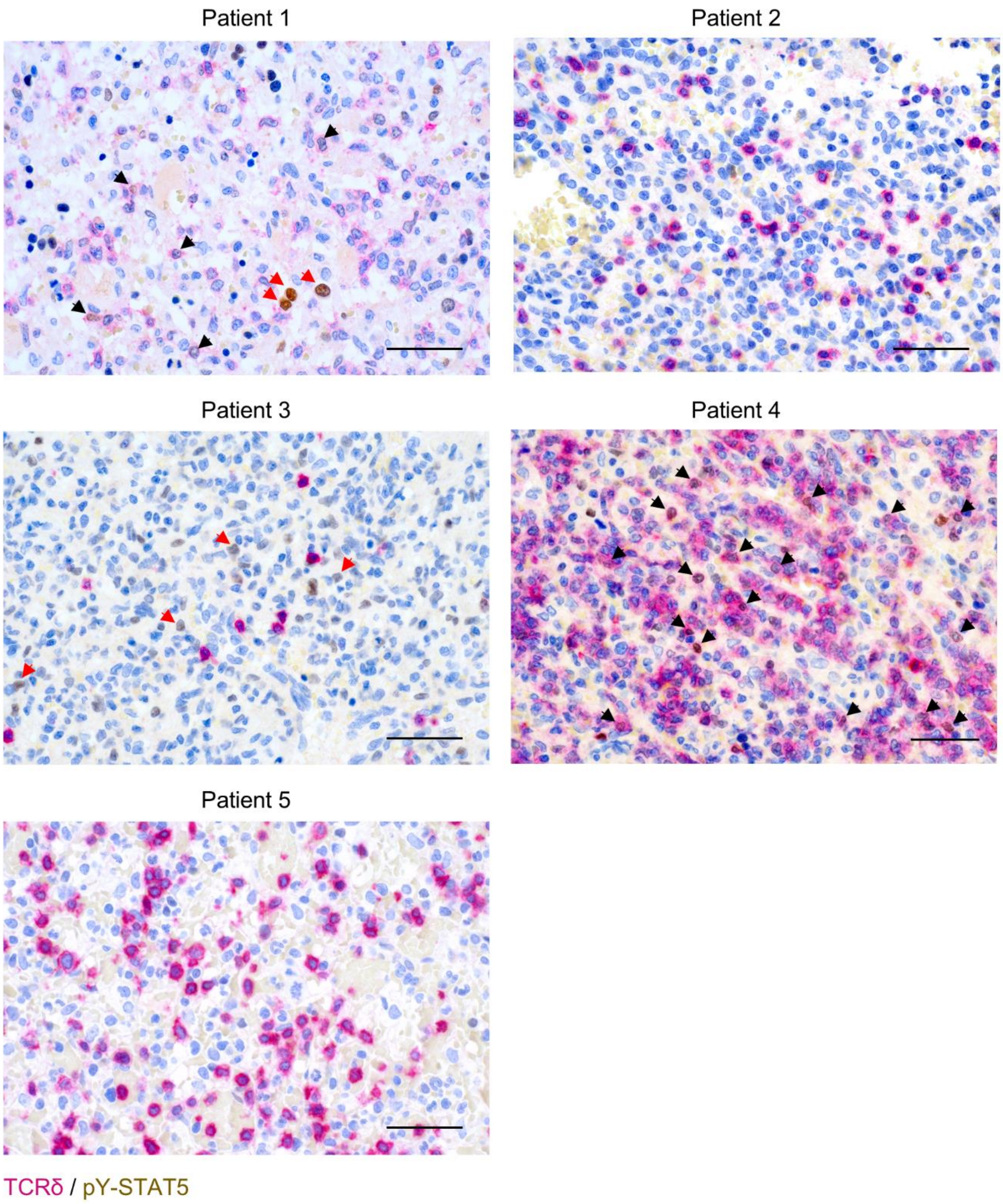


**Supplemental Figure 11.** Representative images from IHC analysis of TCRδ (magenta) and pY-STAT5 (brown) double staining in the bone marrow (patient 1) or spleen (patients 2-5) of HSTCL patients, imaged by light microscopy (scale bar = 50 μm). Black arrows indicate tumor cells with TCRδ and pY-STAT5 double staining; red arrows indicate positive pY-STAT5 staining in surrounding, non-malignant cells (e.g. strong nuclear staining in erythroid precursors, patient 1; weak nuclear staining in reactive lymphoid cells; patient 3).

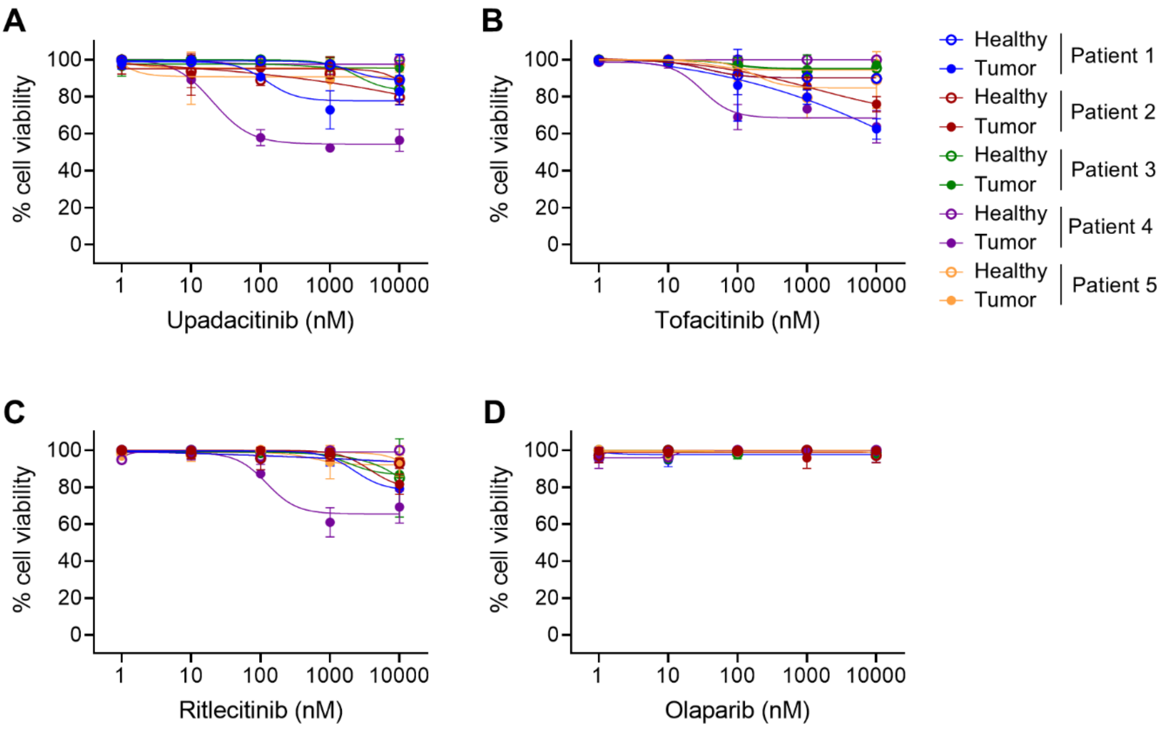


**Supplemental Figure 12. A-D)** Cell viability curves upon 24 hr treatment of A) upadacitinib, B) tofacitinib, C) ritlecitinib or D) olaparib at various concentrations on primary HSTCL patient samples containing both tumor and healthy cell populations, distinguished by cell surface markers (see Supplementary Table 1) and flow cytometry. Data are graphed as mean (± SD) of technical duplicates from one experiment (*n* = 1).

### Supplementary References


1. Pham HTT, Maurer B, Prchal-Murphy M, et al. STAT5BN642H is a driver mutation for T cell neoplasia. *J Clin Invest*. 2018;128(1):387-401.
2. Picelli S, Faridani OR, Bjorklund AK, Winberg G, Sagasser S, Sandberg R. Full-length RNA-seq from single cells using Smart-seq2. *Nat Protoc*. 2014;9(1):171-181.
3. Schmieder R, Edwards R. Quality control and preprocessing of metagenomic datasets. *Bioinformatics*. 2011;27(6):863-864.
4. Dobin A, Davis CA, Schlesinger F, et al. STAR: ultrafast universal RNA-seq aligner. *Bioinformatics*. 2013;29(1):15-21.
5. Li H, Handsaker B, Wysoker A, et al. The Sequence Alignment/Map format and SAMtools. *Bioinformatics*. 2009;25(16):2078-2079.
6. Danecek P, Bonfield JK, Liddle J, et al. Twelve years of SAMtools and BCFtools. *Gigascience*. 2021;10(2):giab008.
7. Liao Y, Smyth GK, Shi W. featureCounts: an efficient general purpose program for assigning sequence reads to genomic features. *Bioinformatics*. 2014;30(7):923-930.
8. Love MI, Huber W, Anders S. Moderated estimation of fold change and dispersion for RNA-seq data with DESeq2. *Genome Biol*. 2014;15(12):550.
9. Wickham H. Ggplot2: Elegant Graphics for Data Analysis (ed 2nd). New York: Springer; 2016.
10. Carlson M. Orthology.Eg.Db: Orthology Mapping Package; 2024.
11. Carlson M. Org.Mm.Eg.Db: Genome Wide Annotation for Mouse; 2024.
12. Carlson M. Org.Hs.Eg.Db: Genome Wide Annotation for Human. 2024.
13. Finalet Ferreiro J, Rouhigharabaei L, Urbankova H, et al. Integrative genomic and transcriptomic analysis identified candidate genes implicated in the pathogenesis of hepatosplenic T-cell lymphoma. *PLoS One*. 2014;9(7):e102977.
14. Gu Z, Eils R, Schlesner M. Complex heatmaps reveal patterns and correlations in multidimensional genomic data. *Bioinformatics*. 2016;32(18):2847-2849.
15. Subramanian A, Tamayo P, Mootha VK, et al. Gene set enrichment analysis: a knowledge-based approach for interpreting genome-wide expression profiles. *Proc Natl Acad Sci U S A*. 2005;102(43):15545-15550.
16. Nabekura T, Gotthardt D, Niizuma K, et al. Cutting Edge: NKG2D Signaling Enhances NK Cell Responses but Alone Is Insufficient To Drive Expansion during Mouse Cytomegalovirus Infection. *J Immunol*. 2017;199(5):1567-1571.
17. Ye J, Coulouris G, Zaretskaya I, Cutcutache I, Rozen S, Madden TL. Primer-BLAST: A tool to design target-specific primers for polymerase chain reaction. *BMC Bioinformatics*. 2012;13:134.

18. Malkki M, Petersdorf EW. Genotyping of single nucleotide polymorphisms by 5′ nuclease allelic discrimination. *Methods in Molecular Biology*. 2012;882:173-182.
19. Echwald SM, Andreasen D, Mouritzen P. LNA™ Adding New Functionality to PCR. PCR Technology: Current Innovations, Third Edition; 2013:87-102.
20. Wen TT, Zhang XH, Lippuner C, Schiff M, Stuber F. Development and Evaluation of a Droplet Digital PCR Assay for 8p23 β-Defensin Cluster Copy Number Determination. *Molecular Diagnosis & Therapy*. 2021;25(5):607-615.
21. Shin SB, Lo BC, Ghaedi M, et al. Abortive gammadeltaTCR rearrangements suggest ILC2s are derived from T-cell precursors. *Blood Adv*. 2020;4(21):5362-5372.
22. Heilig JS, Tonegawa S. Diversity of murine gamma genes and expression in fetal and adult T lymphocytes. *Nature*. 1986;322(6082):836-840.
